# Pharmacological blockade of opioid receptors in the ventrolateral periaqueductal gray does not affect avoidance acquisition or performance in rats

**DOI:** 10.64898/2025.12.17.692586

**Authors:** Laura Vercammen, Mirte De Ceunink, Tom Beckers, Bram Vervliet, Laura Luyten

## Abstract

Avoidance learning involves associating a behavioral response with the omission of an expected threat, but the neural mechanisms that drive this learning remain unclear. Research on fear extinction points to a critical role for opioid receptors in the ventrolateral periaqueductal gray (vlPAG) in computing the aversive prediction error signal that is generated when there is a difference between expected and actual aversive events. Based on these fear extinction findings, we hypothesized that vlPAG opioid signaling might also support the early stages of avoidance learning. To test this, 15 Wistar rats (7 females, 8 males) received intra-vlPAG infusions of either naloxone hydrochloride (n = 7, 2.5 µg/0.5 µl), a non-selective opioid receptor antagonist, or vehicle (n = 8, 0.5 µl), immediately before the first and second session of two-way active avoidance training. No infusions were given before the third session, to examine avoidance performance under continued, drug-free acquisition. For the fourth and final session, drug conditions were reversed to examine the acute effect of naloxone on already established avoidance performance. Our results indicate that unilateral intra-vlPAG naloxone, at the tested dose and timing, did not significantly impair avoidance acquisition, nor its performance. These findings do not support a role for opioid signaling in the vlPAG during the initial learning or expression of two-way active avoidance.

## Introduction

Actively learning to recognize cues that predict danger is critical for survival across species (Rochais et al., 2022). In rodents, for example, the ability to associate specific environmental cues (e.g., odors or sounds) with the presence of a predator enables the implementation of adaptive defensive strategies to increase the likelihood of survival (Ferrero et al., 2011). One such strategy is active avoidance, where animals learn to associate a behavioral response with the omission of the expected threat (Manning et al., 2020). For instance, a wild rat foraging along a hedge may learn that upon hearing the screech of an approaching hawk, rapidly crossing to the opposite side of the hedge prevents an attack. While the neural circuits that guide the formation of cue-threat associations are well established (e.g., Maren & Quirk, 2004; Fanselow & Poulos, 2005; Sigurdsson et al., 2007; Medina et al., 2002), understanding the neural systems that drive active avoidance defensive strategies proves to be more challenging, as avoidance is learned and reinforced in the absence of an explicit observable outcome (i.e., an avoidance response omits the threat from occurring; Cain, 2019).

In the laboratory, avoidance learning can be modeled using the two-way active avoidance (2WAA) task (Fernández-Teruel & Tobeña, 2020; LeDoux et al., 2017). In this task, rats are presented with repeated pairings of a conditioned stimulus (CS, e.g., a tone) with an aversive unconditioned stimulus (US, e.g., a foot shock). Consequently, rats learn to anticipate the US whenever the CS is presented (Cain, 2019). With repeated trials, rats learn that they can avoid receiving the foot shock US by shuttling to the opposite compartment of the shuttle box during CS presentation.

Avoidance learning is thought to be reinforced by the expectancy violation, or prediction error (PE), that accompanies it (Moutoussis et al., 2008). While rats initially expect to receive the aversive US at the end of the CS, shuttling to the opposite compartment effectively omits the foot shock, creating a mismatch between expectations (i.e., a foot shock at the end of the CS) and the actual outcome (i.e., no shock and safety instead). Findings from Pavlovian fear extinction research, which involves CS-alone presentations following cued fear conditioning with the aim of reducing CS-elicited fear, suggest that such discrepancies between an expected aversive US and the lack of actual US received produce an aversive prediction error (PE) signal that is critical in guiding extinction learning (McNally et al., 2011). Specifically, whenever an aversive US exceeds expectations (in the case of fear conditioning) or is less than expected (in the case of extinction learning), a neural PE signal is generated that serves as a teaching signal to update threat expectancies, by regulating synaptic plasticity of lateral amygdala (LA) neurons, where CS-threat associations are stored and updated to guide CS-related learning (LeDoux et al., 2000).

The aversive PE teaching signal is thought to originate from the ventrolateral region of the midbrain periaqueductal gray (vlPAG). Over the course of fear conditioning, US-evoked responding in LA and PAG neurons declines as the US becomes more expected, and pharmacological inactivation of the vlPAG reduces US-evoked responding in LA neurons (Johansen et al., 2010), in line with the notion that an aversive PE signal arises from the vlPAG and influences LA synaptic plasticity. Furthermore, the nociceptive ascending pathways that convey information about the actual US on a conditioning trial and the cortical projections that carry information about US expectancies converge in the vlPAG (Tracey, 2004; McNally et al., 2011), making it a prime candidate for sending an instructive aversive teaching signal to modulate amygdala synaptic plasticity and update threat expectancies to guide new learning.

There is evidence that this PE signal is regulated by endogenous opioids in the vlPAG. Pharmacological blocking of opioid receptors by either systemic (McNally et al., 2003; Kim & Richardson, 2009) or intra-vlPAG (McNally et al., 2004b; Parsons et al., 2010) administration of the non-selective opioid receptor antagonist naloxone significantly impairs fear extinction learning, presumably by reducing the discrepancy between the actual and the expected outcome of a conditioning trial, resulting in a smaller PE and less extinction learning (McNally et al., 2011). Interestingly, this effect seems to be driven by the actions of endogenous opioids at µ-opioid receptors (MORs), a specific subtype of opioid receptors. Naloxone has the highest affinity for the µ-opioid receptor at the doses that were used for intra-vlPAG infusions (Mansour et al., 1995), and at the cellular level, MORs have the highest contribution to the actions of vlPAG opioids (Chieng & Christie, 1994). Indeed, administration of antagonists that act on δ-opioid receptors (e.g., naltrindole) or κ-opioid receptors (e.g., nor-BNI) in the vlPAG does not impair fear extinction learning, suggesting a primary role for MORs in mediating the aversive PE that is necessary for fear extinction learning (McNally et al., 2005).

A more direct role for endogenous opioids acting at MORs in the vlPAG in encoding the aversive PE during fear learning comes from studies that evaluated the effect of opioid antagonists on the blocking of fear conditioning. In blocking paradigms, rats are trained to fear a visual CS by presenting it with an aversive US (e.g., a foot shock) in stage 1. In stage 2, rats receive compound presentations of the visual CS with an auditory CS followed by the US. Fear learning to the auditory CS is blocked during stage 2, because the actual outcome of the trial is not different from the expected outcome (i.e., there is no PE; Kamin, 1968; but see Maes et al., 2016). Using a within-subjects design that also included a control CS which was not conditioned during stage 1, but was presented in compound during stage 2 (and hence, based on a PE account, would enhance fear when later tested), McNally & Cole (2006) showed that systemic naloxone impaired blocking, but did not affect fear learning to the control CS, supporting the role of opioid receptors in mediating the aversive PE directly. Again, this effect was dependent on MORs, as infusions of a specific µ-opioid receptor antagonist (CTAP) into the vlPAG similarly prevented the blocking of fear conditioning.

Together, these findings suggest that µ-opioid receptors in the vlPAG play a critical role in the generation of an aversive PE signal that conveys information about the discrepancy of expected and actual aversive events, which is critical for both fear and extinction learning. We propose that a similar aversive PE signal might underly avoidance acquisition, as avoidance learning similarly involves learning through the absence of an expected threat (Moutoussis et al., 2008; Manning et al., 2020). While initial evidence supports the involvement of opioids in avoidance, as systemic naloxone administration has been found to impair its acquisition (Bennet & Hock, 1990; Turnbull et al., 1983), the potential mediating role of the vlPAG has not yet been examined. In this study, we evaluated whether the non-selective opioid receptor antagonist naloxone administered into the vlPAG impairs the acquisition of avoidance, by attenuating the aversive PE signal that occurs when aversive events are unexpectedly omitted.

## Methods and Materials

### Preregistration and data availability

The experiment was preregistered on the Open Science Framework (OSF) (https://osf.io/w3256/). The raw data files, processed data files and R scripts generated for the current study are available via the same link. Video files are available upon request.

### Subjects

Ten male (weighing between 250 – 275 grams upon arrival) and 10 female (weighing between 225 – 250 grams upon arrival) Wistar rats were purchased from Janvier Labs (Le Genest Saint Isle, France). Upon arrival in the lab, rats were housed individually in standard animal cages with wooden sticks as cage enrichment and kept under conventional laboratory conditions (12-h day-night cycle; lights on 7:00-19:00; 22°C) with ad libitum access to food and water. Experimental sessions were conducted between 8:00 and 18:00. The number of animals per group was calculated using effect sizes found in the literature (*f* > 0.40, adjusted to *f* = 0.40) to achieve a power of .80, accounting for an expected exclusion of up to 4 animals (G-Power version 3.1). Both sexes were included in this study for the sake of representativeness (Dalla et al., 2024). Rats were handled briefly for 4 days prior to behavioral testing. All experiments were approved by the KU Leuven animal ethics committee (project P027/2025) and conducted in accordance with the Belgian and European laws, guidelines and policies for animal experimentation, housing and care (Belgian Royal Decree of 29 May 2013 and European Directive 2010/63/EU on the protection of animals used for scientific purposes of 20 October 2010).

### Stereotaxic surgery

Rats were anesthetized through intraperitoneal injection of a mixture of Ketamine (Nimatek, 100 mg/ml) and Metodomine hydrochloride (Domitor, 1 mg/ml), which was administered at a volume of 200 µl/100 gram. Rats were placed in the stereotaxic frame on a heating pad and given Meloxicam (Metacam, 5 mg/ml; administered at 100 µl/100 g) before a 26-gauge guide cannula (Protech International, Texas, USA; C315G/SPC) was implanted, positioning the tip of the guide cannula 1 mm above the right vlPAG (coordinates according to Paxinos & Watson (1982): AP: -7.56 mm, ML: 0.9 mm, DV: 4.3 mm). The cannula was implanted unilaterally in the right vlPAG to facilitate comparison with prior research (McNally et al., 2004b). The guide cannula was fixed with UV-cured dental cement and anchored to the skull using bone screws (Fine Science Tools, Heidelberg, Germany). A dummy (Protech International, C315DC/SPC) was inserted into the guide cannula and kept in place at all times, except during infusions. Immediately after surgery, rats were subcutaneously injected with 2.5 ml 0.9% physiological saline for rehydration. Twenty-four hours after surgery, rats were again given Meloxicam (Metacam, 5 mg/ml; administered at 100 µl/ 100 g). Rats were allowed to recover for 6 to 7 days before the start of behavioral procedures.

### Apparatus

Two-way active avoidance training took place in two identical rectangular shuttle boxes (50.8 cm x 25.4 cm x 30.5 cm), each housed inside larger sound-attenuating cubicles (72.4 cm x 41.1 cm x 43.2 cm, Coulbourn Instruments, Pennsylvania, USA). The side walls and ceilings of the shuttle boxes were made of metal, whereas the front and back walls were clear Plexiglas. The shuttle box was divided into two equal compartments by a metal divider placed halfway along the length of the shuttle box (Coulbourn Instruments). Rats could freely move from one side of the box to the other via a passage (8 cm width) in the divider. The floor of the shuttle box was a stainless steel grid, through which scrambled foot shocks (0.4 mA) could be delivered that served as the unconditioned stimulus (US). In each compartment of the shuttle box, a speaker was mounted to the side wall to deliver a 3-kHz tone (presented at +/- 75 dB) that served as the conditioned stimulus (CS). The shuttle box was unilluminated. Two photocell sensor bars, one in each side of the shuttle box, ensured automatic detection of the position of the rat throughout the experiment. All behavioral sessions were videorecorded via an infrared camera (HD IP camera, Foscam C1, Shenzhen, China) mounted to the ceiling of the isolation cubicle.

The first behavioral phase of the experiment (Pavlovian fear conditioning) took place in the same shuttle boxes as used for two-way active avoidance training, except that the passage that divides both compartments was closed off by a metal door that prevented the rats from entering the adjacent compartment.

### Drug administration

Rats were administered either naloxone hydrochloride (2.5 µg/0.5 µl; n = 10; Tocris Bioscience, Bristol, UK) or 0.9% physiological saline (0.5 µl; n = 10) by removing the dummy and inserting a 33-gauge injection cannula with a 1-mm projection (C315I/SPC, Protech International) into the guide (McNally et al., 2004b). Naloxone or vehicle was infused using an infuse/withdraw pump (Pump 11 Pico Plus Elite, Harvard Apparatus, Massachusetts, USA) which contained a 50-µl glass syringe (Hamilton 80521, 705SN 22 gauge, Giarmata, Romania) that was connected to the injection cannula using FEP tubing (0.12 mm inner diameter) and tubing adaptors (Microbiotech/se, Stockholm, Sweden). The drug or vehicle was infused at a flow rate of 0.25 µl/min over a period of 2 minutes, after which the injection cannula was left in place for an additional minute before it was removed and the dummy was reinserted. Immediately after infusion, that rat was placed in the shuttle box (McNally et al., 2004b).

### Behavioral procedures

For an overview of the experimental procedures, see Figure 1.

**Figure 1.**
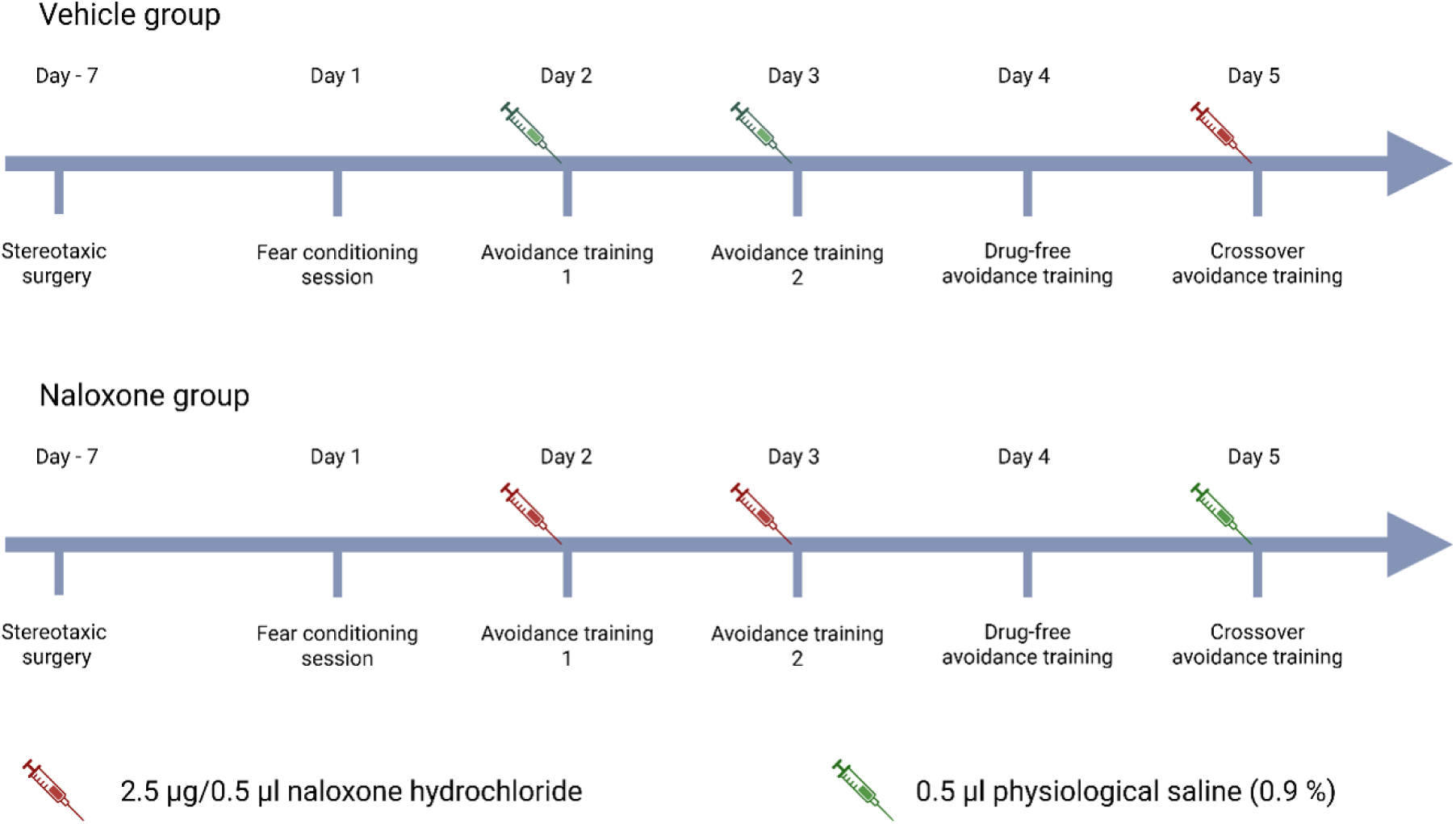
Overview of experimental procedures. Rats underwent stereotaxic surgery followed by a recovery period of 6 to 7 days. The behavioral procedure started with a Pavlovian fear conditioning session in one compartment of the shuttle box, followed by four training sessions in the 2WAA task. Before the first and second avoidance training session, rats received unilateral intra-vlPAG infusions of either naloxone (2.5 µg/0.5 µl; final n = 7) or vehicle (0.5 µl; final n = 8). No infusions were given before the third, drug-free, training session. For the final, crossover, training session, drug administration was reversed between the two groups.

#### Pavlovian fear conditioning

The behavioral procedure began with a Pavlovian fear conditioning session to ensure that all subjects acquired the CS–US association prior to the onset of avoidance training. This step was necessary to standardize threat learning across groups, and to keep it drug-free and separate from the subsequent instrumental learning phase. Successful avoidance learning depends on the prior formation of a CS–threat association (LeDoux et al., 2017; Mowrer & Lamoreaux, 1946), and previous studies have shown that intra-vlPAG administration of naloxone enhances such associative learning (McNally et al., 2004a). Therefore, we separated the Pavlovian and instrumental components of the task. Specifically, the CS–US association was established before introducing the opportunity to perform an avoidance response, thereby excluding any effects (e.g., mediated by hypoalgesia) of opioid antagonism on the acquisition of the CS-US association. This allowed us to isolate the effect of interest, namely the prediction-error-related effects of naloxone on avoidance learning, when the to-be-learned response is characterized by US omission.

For Pavlovian fear conditioning, the rats were introduced to a single compartment of the shuttle box (side counterbalanced between rats). After five minutes of habituation, the rats were given 5 presentations of the CS (3-kHz tone, 20 s, +/- 75 dB) immediately followed by a 1-second US (0.4-mA foot shock). The intertrial interval (ITI) averaged around 3 minutes (60 s – 260 s). Sixty seconds after the final CS-US presentation, rats were removed from the setup.

#### Two-way active avoidance training

Twenty-four hours after the fear conditioning session, the rats were subjected to four avoidance training sessions in the shuttle boxes, that took place on four consecutive days. Immediately before the first and second avoidance training session, rats were infused with either naloxone or vehicle. The third avoidance training session took place in the absence of the drug or vehicle (i.e., the drug-free avoidance training session). Finally, for the fourth avoidance training session, drug conditions were reversed (i.e., the crossover avoidance training session). Specifically, rats of the naloxone group received the vehicle before the fourth session, whereas rats of the vehicle group received naloxone, enabling us to examine acute effects of naloxone on established avoidance performance. Each avoidance training session started with a 5-minute acclimation period during which no stimuli were delivered. After acclimation, the rats were given 30 presentations of the CS and US, with an ITI averaging around 1 minute (30 s – 90 s). The CS lasted a maximum of 20 seconds and was followed immediately by the US which lasted a maximum of 10 seconds. If the rat shuttled to the other compartment during CS presentation, the CS was terminated and no shock was given, i.e., the US was avoided and the trial was labeled as an avoidance response. If the rat crossed to the other compartment during the presentation of the US, the shock was terminated and the trial was labeled as an escape response. The total duration of the training session was 35 minutes. The number of avoidance responses, escape responses and escape failures (i.e., the rat remained in the compartment throughout the 10-s US presentation) were recorded via Graphic State 4 software (Coulbourn Instruments), which also controlled the delivery of the stimuli. Freezing, i.e., the absence of body movement except for movement related to respiration (Fanselow, 1994), was scored from recorded videos by an experienced observer blinded to session and group.

### Histological analyses

At the end of the behavioral procedures, the animals were given an overdose of sodium pentobarbitol (200 mg/ml solution for injection, Vetoquinol B.V.; 300 mg/kg). Rats were perfused transcardially with 1% phosphate-buffered saline, followed by 4% paraformaldehyde, before removing the brain. The fixed brains were sectioned coronally at 50 µm using a vibratome. Every third section was collected and stained using 0.5% cresyl violet acetate (Carl Roth, Karlsruhe, Germany). The placement of the cannulas was verified using the boundaries defied by Paxinos and Watson (1998; Figure 2). When the cannula tip was outside the vlPAG, the subject was excluded from subsequent analyses. In total, 5 subjects were excluded because of incorrect probe location, resulting in a total sample size of 15 animals, of which 7 were in the naloxone group (3 females and 4 males) and 8 in the vehicle group (4 females and 4 males).

**Figure 2.**
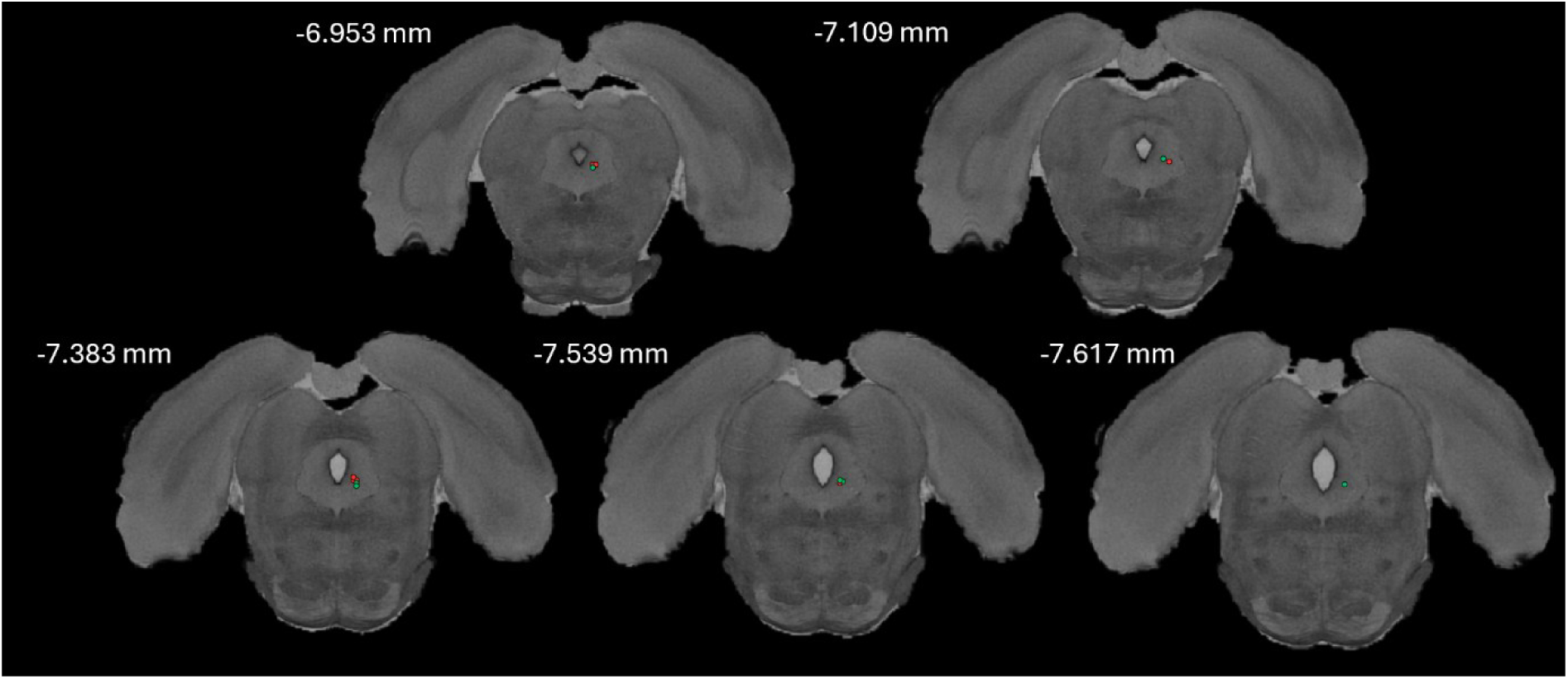
Cannula placement in the vlPAG. Dots indicate the position of the tip of the injection cannula for the 15 included animals. Distance posterior to bregma is indicated for each coronal section. Red: naloxone group, Green: vehicle group. Figure created with Waxholm Space Atlas.

### Statistical analyses

All analyses were preregistered on the Open Science Framework, unless stated otherwise. Data are expressed as means ± SEM and analyzed using R statistical software (RStudio version 3.5.1). The mean number of avoidance responses was calculated per block of 10 trials within each avoidance training session and analyzed using a mixed design ANOVA (repeated-measures factor: Block, between-subjects factors: Group and Sex), to evaluate avoidance responding over the course of each avoidance training session (Vercammen et al., 2025). Moreover, an exploratory trial-based analysis was performed for each avoidance training session, by coding the data into a binary format (successful avoidance = 1, escape response/escape failure = 0) and applying a generalized linear mixed-effects model to the binary outcome with fixed-effects parameters Trial, Group and the interaction between Trial and Group, and a random effects parameter Subject to account for within-subject variability due to the repeated measures design, using the R-package lme4 (https://cran.r-project.org/package=lme4). Preregistered secondary analyses on freezing, response latencies, shuttles during the first 5 minutes of habituation and the ITIs of each avoidance training session and escape failures can be found in the Supplemental Material.

### Additional analyses

As an additional control to establish if infusion of 2.5 µg naloxone in the right vlPAG produced any measurable functional effects, we analyzed grooming and shock-elicited freezing, after completion of the analyses described above. These additional analyses were preregistered on OSF (https://osf.io/zryk8/) before manually scoring grooming and shock-elicited freezing from video recordings.

We primarily looked at shock-elicited freezing and did so early on during avoidance training, when the animals had not yet acquired the avoidance response very well and still received a considerable amount of shocks. Prior research in female and male rats (Fanselow & Bolles, 1979; Hammer & Kapp, 1986) has shown that naloxone (administered either systemically or unilaterally into the vlPAG, albeit at more anterior coordinates) increases shock-elicited freezing, and may also decrease grooming (defined as grooming, scratching and licking the body). Fanselow & Bolles (1979) also found effects of naloxone on grooming in non-shocked animals. As our behavioral procedure was quite different from the ones used in prior research, we focused our analysis on shock-elicited reactions early on, during the first block of avoidance training 1 (i.e., the first 10 trials).

Specifically, we analyzed freezing and grooming (both expressed as percentage of time) during the 30 seconds following shock delivery (i.e., on non-avoided trials during the first block of avoidance training 1). Freezing and grooming were scored by an experimenter blinded to group allocation. The measurement after the first shock was excluded, in line with observations by Fanselow & Bolles (1979). This means that animals that avoided 9 or 10 trials of the first block were excluded from the shock-elicited behavior analyses. For each animal, the average percentage of freezing and grooming was taken for all non-avoided trials (except the first one) during block 1. In addition, we measured grooming during the first 5 minutes of acclimation (before any tones or shocks were given) in all animals.

Our primary hypothesis was that naloxone animals would freeze more than saline animals during the post-shock period. This was assessed with a non-parametric one-sided Mann-Whitney U test.

We had two secondary hypotheses related to grooming. The first one was that naloxone animals would show less grooming than saline animals during the post-shock period. The planned one-sided t-test was not conducted because the animals did not show any grooming during this period. The second secondary hypothesis was that naloxone animals would show less grooming than saline animals during the 5-min acclimation period of avoidance training session 1. This was assessed with a one-sided t-test.

The results of these additional analyses (see Supplemental Material) did not alter any of the conclusions drawn from the main analyses. They failed to provide a positive control demonstrating that intra-vlPAG naloxone produced a measurable behavioral effect in our rats.

## Results

For the sake of brevity, only the main statistical results are reported. For a detailed overview of all statistical results, see Supplementary Tables 1 – 4. Graphs that display the block data for each sex separately are included as a Supplementary Figure (Suppl. Fig. 6).

### Avoidance training under influence of naloxone or vehicle

To evaluate whether naloxone impaired avoidance acquisition, a mixed ANOVA with repeated-measures factor Session (2 levels: session 1 and 2) and between-subjects factors Group (2 levels: naloxone and vehicle) and Sex (2 levels: male and female) was conducted (Figure 3A). The number of avoidance responses significantly increased from session 1 to session 2 (*F*(1,11) = 17.547, *p* = .002, ƞ*_p_*² = 0.615). However, no significant effects of Group (*F*(1,11) = 0.099, *p* = .759, ƞ*_p_*² = 0.009) or Sex (*F*(1,11) = 1.547, *p* = .239, ƞ*_p_*² = 0.123) were observed. There was also no significant Group by Session interaction (*F*(1,11) = 0.942, *p* = .353, ƞ*_p_*² = 0.079), nor any significant interactions with Sex (see Suppl. Table 1). These findings suggest that naloxone did not impair avoidance acquisition, compared to vehicle-administered rats.

**Figure 3.**
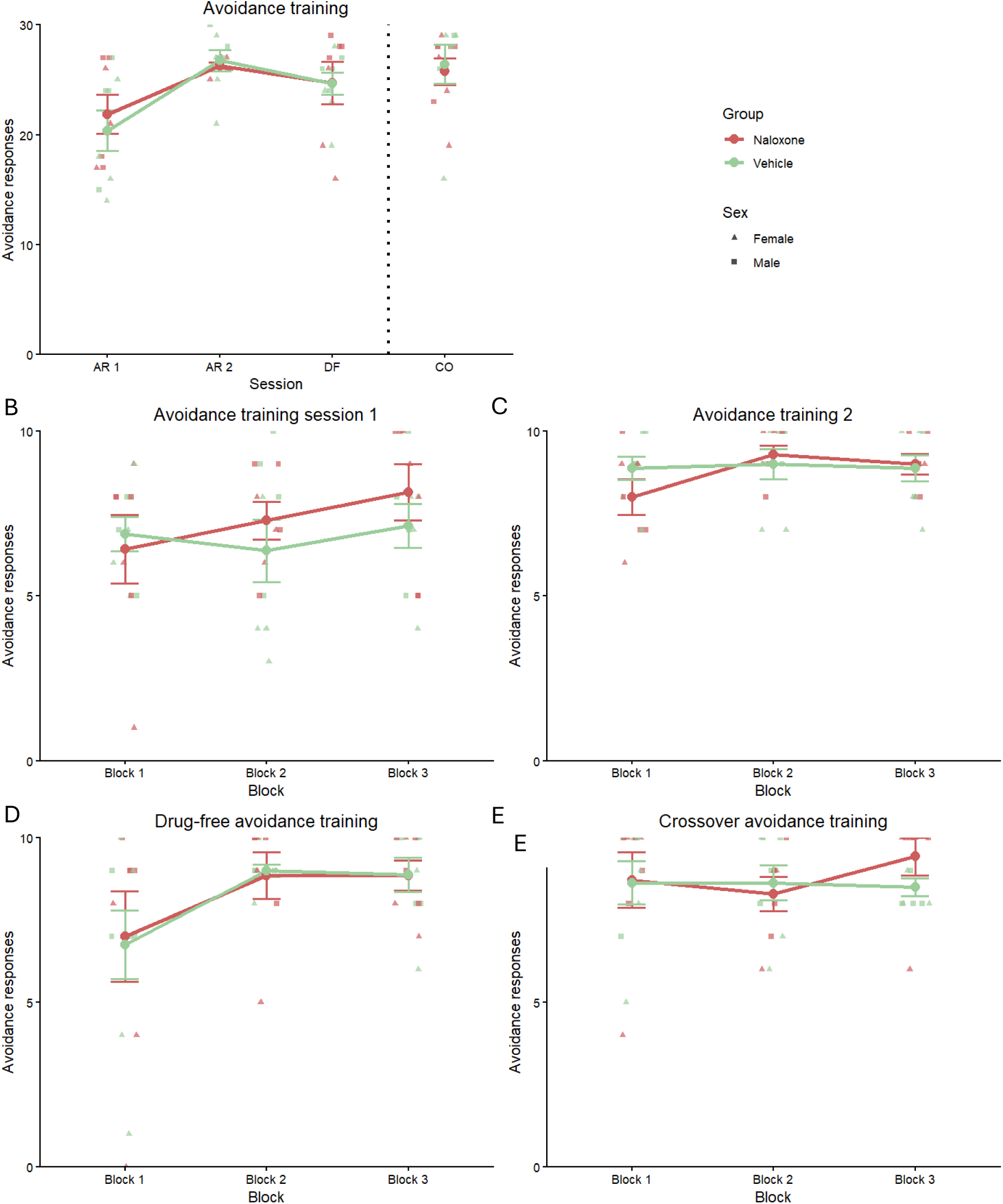
Two-way active avoidance training data. (A) Mean (± SEM) number of avoidance responses for each avoidance training session separately for the naloxone (n = 7; Red) and vehicle (n = 8; Green) groups. (B) Mean (± SEM) number of avoidance responses in three blocks of 10 trials each for rats that received naloxone (n = 7) and vehicle (n = 8) prior to avoidance training session 1. (C) Mean (± SEM) number of avoidance responses in three blocks for rats that received naloxone (n = 7) and vehicle (n = 8) prior to avoidance training session 2. (D) Mean (± SEM) number of avoidance responses in three blocks of 10 trials each for rats that previously received naloxone (n = 7) and vehicle (n = 8) prior to avoidance training session 1 and 2 (i.e., the drug-free avoidance training session). (E) Mean (± SEM) number of avoidance responses in three blocks of 10 trials each for rats of the naloxone (n = 7) and vehicle (n = 8) groups during the crossover avoidance training session. During this session rats from the vehicle group (green) received an infusion of naloxone, whereas rats of the naloxone group (red) received a vehicle infusion. Triangles depict individual datapoints for female subjects and squares depict individual datapoints for male subjects. See Suppl. Fig. 5 for an overview of the block data for each avoidance training session plotted for each sex separately.

This was further corroborated when zooming into the data of the first avoidance training session by splitting the data into three blocks of 10 trials each. We did not observe any significant effects of Group (*F*(1,11) = 0.290, *p* = .601, ƞ*_p_*² = 0.026), Sex (*F*(1,11) = 0.964, *p* = .347, ƞ*_p_*² = 0.081) or Block (*F*(10.03,52.94) = 2.084, *p* = .168, ƞ*_p_*² = 0.159; Figure 3B). Moreover, there was no significant Group by Block interaction effect (*F*(6.29,52.94) = 1.306, *p* = .286, ƞ*_p_*² = 0.106), nor any significant interaction effects with Sex (Suppl. Table 2). Finally, we conducted a generalized linear mixed model (GLMM) analysis to examine the effects of Trial and Group (reference group: vehicle) on the binary avoidance data (avoidance = 1, escape/escape failure = 0), with a random intercept for each subject. The logistic regression coefficients indicated no significant effects of Trial (*β* = 0.016, *SE* = 0.017, *Z* = 1.003, *p* = .316) or Group (*β* = 0.058, *SE* = 0.561, *Z* = 0.104, *p* = .917), nor a significant interaction between Trial and Group (*β* = 0.021, *SE* = 0.025, *Z* = 0.832, *p* = .406; Figure 4A). These findings indicate that there was no significant difference in avoidance responses throughout the first avoidance training session between naloxone-administered rats and vehicle-administered rats.

**Figure 4.**
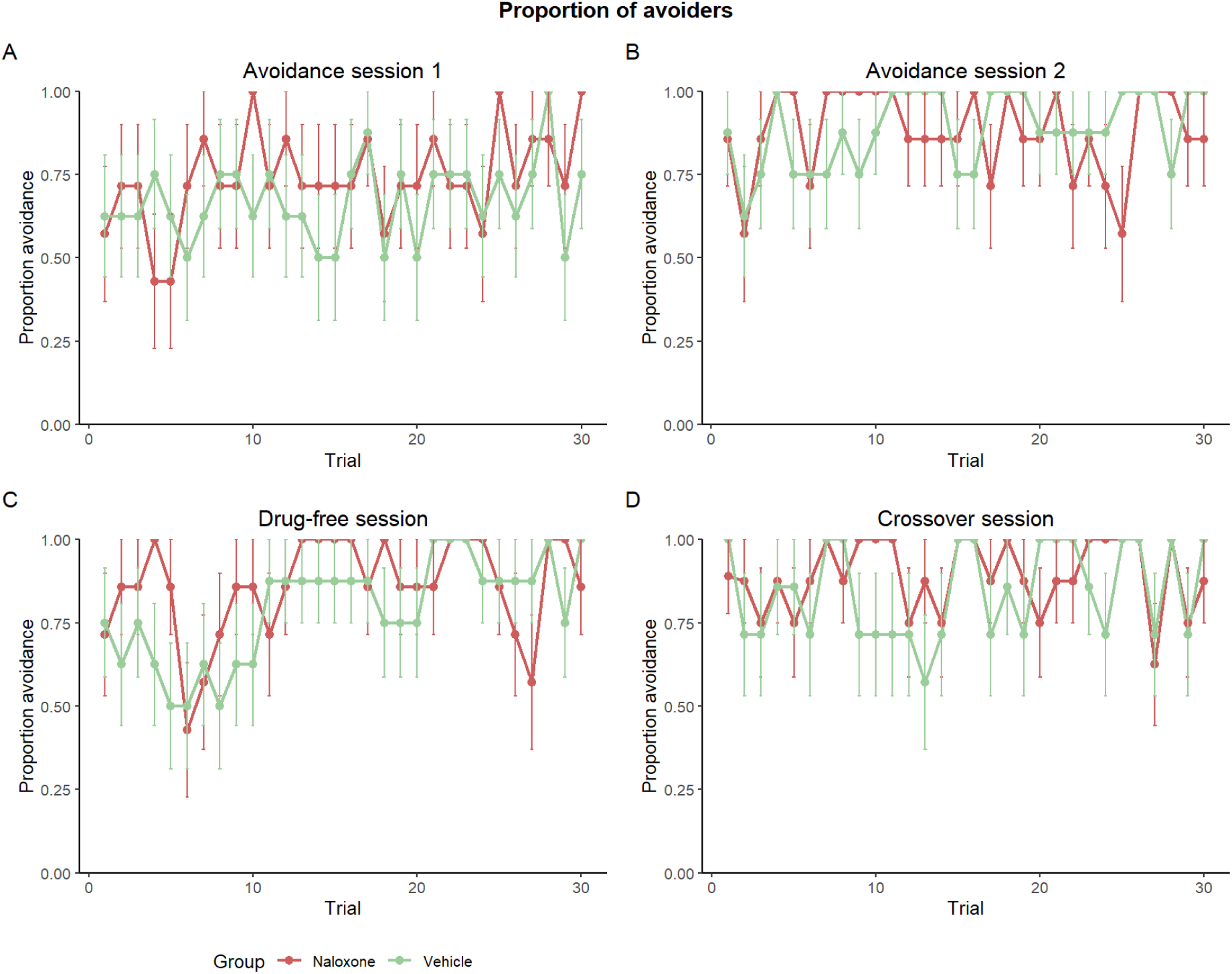
Proportion of avoiders per trial ± SEM for the naloxone (n = 7; red) and vehicle (n = 8; green) groups. for (A) avoidance training session 1, (B) avoidance training session 2, (C) the drug-free avoidance training session, and (D) the crossover avoidance training session.

Similarly, during the second avoidance training session, we observed no significant effects of Group (*F*(1,11) = 0.192, *p* = .670, ƞ*_p_*² = 0.017), Sex (*F*(1,11) = 1.335, *p* = .272, ƞ*_p_*² = 0.108), or Block (*F*(2,22) = 1.731, *p* = .200, ƞ*_p_*² = 0.136). Moreover, there was no significant Group by Block interaction effect (*F*(2,22) = 1.282, *p* = .297, ƞ*_p_*² = 0.104; Figure 3C). The interactions with Sex were also non-significant (Suppl. Table 2). Finally, a generalized linear mixed model analysis indicated a significant ^A^ t of Trial (*β* = 0.062, *SE* = 0.025, *Z* = 2.442, *p* = .015), suggesting ongoing learning, but no effect of Group (*β* = 0.899, *SE* = 0.602, *Z* = 1.495, *p* = .135; Figure 4B). While the interaction between Group and Trial approached significance (*β* = -0.069, *SE* = 0.035, *Z* = -1.934, *p* = .053), suggesting a tendency for naloxone to impair avoidance acquisition over the course of the second training session, this trend disappeared when the data were analyzed in blocks.

### Drug-free avoidance training session

When the rats were tested the following day, without naloxone or vehicle administered immediately before the session, we did not observe a significant effect of Group (*F*(1,11) = 0.085, *p* = .776, ƞ*_p_*² = 0.008; Figure 3D), suggesting that they acquired the avoidance response to a similar extent, regardless of their drug treatment on the previous days. Interestingly, we did observe a significant main effect of Sex (*F*(1,11) = 9.944, *p* = .009, ƞ*_p_*² = 0.475), with females performing fewer avoidance responses than males (*t_Tukey_* = -3.153, *p* = .009), but no significant interaction effects with Sex (Suppl. Table 2). We also observed a significant effect of Block (*F*(1.29,14.17) = 5.246, *p* = .031, ƞ*_p_*² = 0.323; Figure 3D), with rats significantly increasing their performance from block 1 to block 2 (*t*_Tukey_ = -2.832, *p* = .029). Finally, a generalized linear mixed model analysis indicated a significant effect of Trial (*β* = 0.091, *SE* = 0.022, *Z* = 4.121, *p* < .001), in line with the block data, but no effect of Group (*β* = 1.021, *SE* = 0.626, *Z* = 1.630, *p* = .103), nor a significant interaction between Trial and Group (*β* = -0.051, *SE* = 0.032, *Z* = -1.570, *p* = .116; Figure 4C). Together, these findings indicate that there was no significant difference in avoidance responding when rats were previously given naloxone or vehicle, suggesting that both groups acquired the avoidance response similarly, and continued to do so during this session.

### Crossover avoidance training session

Immediately before the crossover training session, rats that previously received naloxone received a vehicle, whereas the control group received naloxone into the vlPAG, enabling us to examine the acute effect of naloxone on avoidance performance, excluding any context effects that could arise because of prior exposure to naloxone. We did not observe an acute effect of naloxone on avoidance responding, as illustrated by a nonsignificant effect of Group (*F*(1,11) = 0.044, *p* = .837, ƞ*_p_*² = 0.004). Moreover, we did not observe significant effects of Sex (*F*(1,11) = 1.360, *p* = 0.268, ƞ*_p_*² = 0.110) or Block (*F*(2,22) = 0.897, *p* = .422, ƞ*_p_*² = 0.075), nor a significant Group by Block interaction effect (*F*(2,22) = 1.472, *p* = .251, ƞ*_p_*² = 0.118; Figure 3E). There were also no significant interaction effects with Sex (Suppl. Table 2). Finally, a generalized linear mixed model analysis indicated no significant effects of Trial (*β* = 0.032, *SE* = 0.023, *Z* = 1.378, *p* = .168) or Group (*β* = 0.991, *SE* = 0.739, *Z* = 1.341, *p* = .180), nor a significant interaction between Trial and Group (*β* = -0.024, *SE* = 0.034, *Z* = -0.716, *p* = .474; Figure 4D). Together, these findings suggest that naloxone did not acutely impair established avoidance performance, as there was no significant difference in performance between naloxone-administered and vehicle-administered rats.

## Discussion

The current study aimed to evaluate the effect of intra-vlPAG administration of naloxone on the acquisition and expression of avoidance. Prior research on fear extinction has identified a critical role for endogenous opioids acting at the µ-opioid receptors (MORs) in the vlPAG in coding the aversive PE signal that updates threat expectancies and controls how much fear and extinction learning takes place (McNally et al., 2009; McNally et al., 2011). Specifically, when naloxone, a non-selective opioid antagonist, was administered intra-vlPAG, extinction learning was significantly impaired (McNally et al., 2004b). As avoidance behavior is thought to be reinforced by the unexpected omission of an aversive event (Moutoussis et al., 2008; Kim et al., 2006), the same opioid-dependent system might track the aversive PE during early avoidance learning. To test this, fifteen Wistar rats (n = 7 females, n = 8 males) were administered either naloxone hydrochloride (n = 7, 2.5 µg/0.5 µl) or vehicle (n = 8, 0.5 µl) before avoidance training. We observed no significant impairment in the acquisition of avoidance following unilateral intra-vlPAG naloxone administration and thus found no evidence that an opioid-dependent mechanism in the vlPAG guides avoidance acquisition, at the tested dose and timing.

Our findings diverge from those in fear extinction research, where intra-vlPAG naloxone administration significantly impaired extinction learning (McNally et al., 2004b). We used the same dose as McNally and colleagues (2.5 µg/0.5 µl), and the same rat strain (Wistar), to facilitate comparisons between studies, with the exception that half of the subjects in our experiment were female. In the same study, a dose-response analysis showed that a higher dose of naloxone (up to 5 µg) resulted in a stronger impairment of extinction learning, as evidenced by more freezing during the subsequent drug-free test (McNally et al., 2004b). It could be that in the context of avoidance learning, a higher dose of naloxone or bilateral infusion would be more effective in impairing avoidance acquisition. Future research may benefit from conducting a dose-response analysis specifically within the framework of avoidance learning.

In our study, naloxone did not produce an increase in freezing per se, nor did it alter nociceptive sensitivity, as indicated by comparable latencies to escape the US between vehicle- and naloxone-administered rats (see Supplemental Material). These observations are consistent with the findings of McNally et al. (2004b), who reported that naloxone neither enhanced freezing to an already extinguished CS, nor affected any component of the aversive foot shock US. Consequently, an important limitation of our study is that it lacks a reliable positive control to verify that naloxone infusion into the vlPAG could have any behavioral effects in our hands. Note that we added the crossover session as an additional control to detect any unwanted, aspecific effects (e.g., motor impairment, other direct effects on the avoidance response per se, instead of on PE). However, we did not actually expect to observe any effects of naloxone during this session where the animals performed an already fully acquired avoidance response (McNally et al., 2004b). In that sense, the crossover test, and the – albeit expected – lack of naloxone effects there, does not help with the limitation mentioned above. A potential strategy to address this limitation would be to directly replicate the experimental procedures of McNally et al. (2004b) within our own laboratory, thereby providing a benchmark for validating drug efficacy.

While we did not observe a significant effect of intra-vlPAG naloxone on avoidance acquisition, previous studies did observe a significant impairment in avoidance learning when naloxone was administered systemically (Bennet & Hock, 1990; Turnbull et al., 1983). Of note, given the systemic administration in those studies, opioid receptors were affected in a brain-wide manner, so the observed effect on avoidance learning may well have been due to the blockade of opioid receptors in other areas than vlPAG. The dorsal PAG (dPAG), for example, seems to be critically involved in controlling escape behaviors and active defensive behaviors such as running and jumping (Lefler et al., 2020; Wendt et al., 2017; Bandler et al., 1988; Fanselow et al., 1995). Dorsal PAG neurons gate and control the initiation of escape, specifically in situations where there is an imminent threat, and previous studies observed a mediating role for opioid receptors in this process (Twardowschy & Coimbra, 2015). Moreover, a recent study identified a dPAG threat-avoidance ensemble, with neurons that are activated during threat avoidance behaviors (Reis et al., 2021), supporting a role of the dPAG in avoidance. Consequently, systemic naloxone may have blocked active defensive behaviors via inhibition of dPAG opioid receptors, which subsequently prevented rats from learning to avoid. Another possibility is that, in prior studies, the systemic administration of naloxone, which is a non-selective opioid antagonist, impaired avoidance responding by blocking kappa opioid receptors (KORs) instead of MORs. Activity of KORs in the VTA of rats has been found to play a crucial role in punishment avoidance. Specifically, Robble et al. (2020) observed that pharmacological blockade of KORs within the VTA disrupted learned quinine avoidance. Importantly, this function of KORs to impair learned avoidance is critically different from the function of MORs, as only the latter have been implicated in aversive PE encoding (McNally et al., 2009). Future studies should further address the relative importance of different opioid receptor subtypes on avoidance acquisition and the mediating role of the dPAG in active avoidance learning.

In summary, we found that intra-vlPAG naloxone administration did not impair avoidance acquisition at the tested dose and timing. This does not align well with previous findings where the same dose of unilateral intra-vlPAG naloxone impaired fear extinction learning (McNally et al., 2004b) as well as with studies reporting impaired avoidance learning following systemic administration of naloxone (Bennet & Hock, 1990; Turnbull et al., 1983). In addition, we found that intra-vlPAG naloxone did not impair the performance of a learned avoidance response. It should be noted that the lack of a positive control demonstrating a measurable behavioral effect of intra-vlPAG naloxone is a major limitation of our study. Keeping this important caveat in mind, we conclude that our findings do not provide support for a role of opioid receptors in the right vlPAG in mediating an aversive PE signal that guides avoidance acquisition.

## Author contributions

Laura Vercammen: conceptualization, data curation, formal analysis, funding acquisition, investigation, methodology, visualization, writing – original draft, writing – review and editing; Mirte De Ceuninck: formal analysis, investigation, writing – review and editing; Tom Beckers: conceptualization, methodology, supervision, writing – review and editing, funding acquisition; Bram Vervliet: conceptualization, methodology, supervision, writing – review and editing, funding acquisition; Laura Luyten: conceptualization, methodology, supervision, writing – review and editing, funding acquisition

## Supplemental Material

For the fear conditioning session, **freezing** during each CS presentation was scored and analyzed using a mixed ANOVA with repeated-measures factor Trial and between-subjects factors Group and Sex. Freezing during each avoidance training was scored during CS 1 and each even CS presentation (i.e., CS 2, CS 4, …, CS 30) and analyzed per session using mixed ANOVAs with repeated-measures factor Trial and between-subjects factor Group and Sex). Moreover, an exploratory analysis on freezing during the first five minutes (habituation) of each avoidance training session was scored and analyzed using a mixed ANOVA with repeated-measures factor Minute and between-subjects factors Group and Sex. The average **latency to avoid** the US, and **escape** the US, were calculated for each avoidance training session, and analyzed using ANOVAs with between-subjects factors Group and Sex. The number of **shuttles** during the 5 minutes (habituation) of each avoidance training session was analyzed using an ANOVA with between-subjects factors Group and Sex. Moreover, the number of shuttles during the ITI of each avoidance training session was analyzed using an ANOVA with between-subjects factors Group and Sex. Finally, we exploratively examined the number of **escape failures**, to evaluate whether naloxone increased failures, compared to vehicle.

## Freezing

### Fear conditioning session

See Supplementary Table 3 for a detailed overview of all statistical results.

**Supplementary Figure 1.**
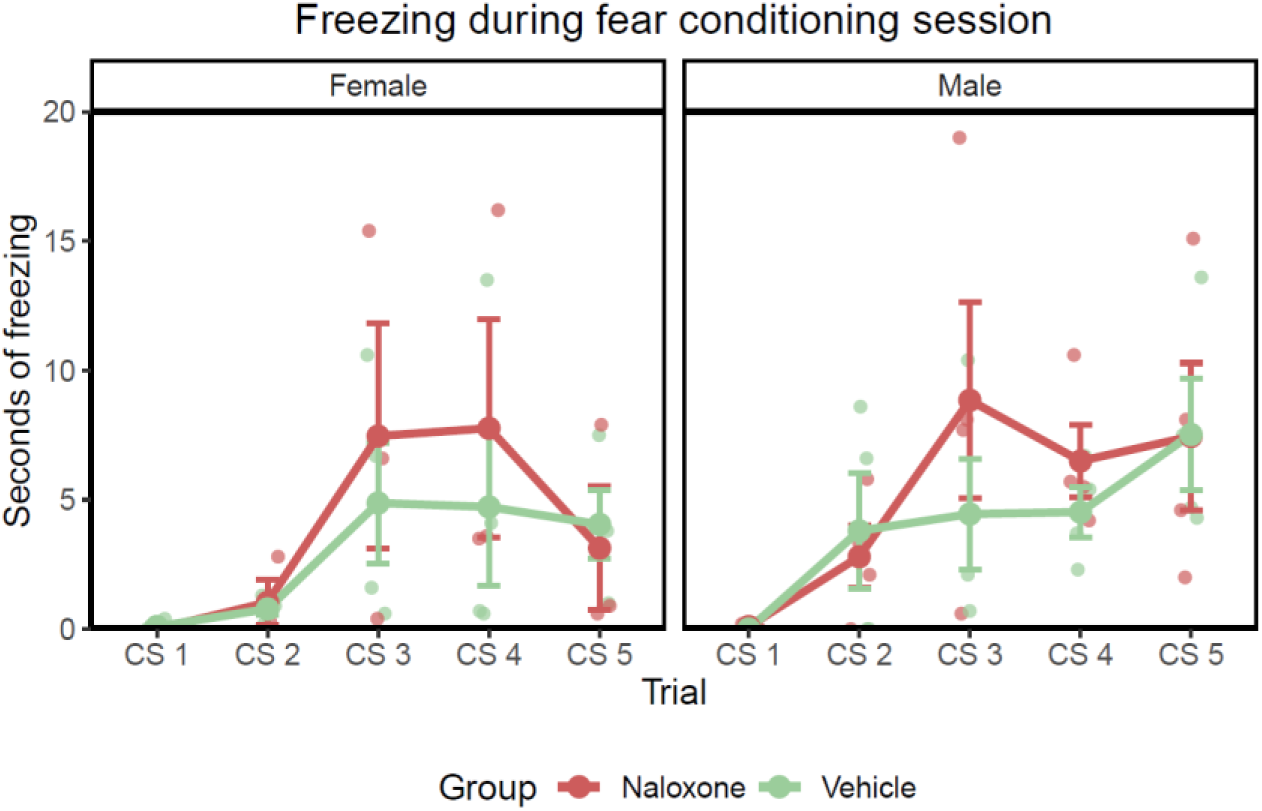
Mean ± SEM freezing during each CS presentation of the fear conditioning session for male and female rats that will either receive a vehicle on the following days, or naloxone.

Freezing during the fear conditioning session was scored during all CS presentations. Freezing significantly increased across the fear conditioning session (*F*(4,44) = 10.034, *p* < .001, ƞ*_p_*² = 0.447). We observed no significant effects of Group (*F*(1,11) = 0.478, *p* = .504, ƞ*_p_*² = 0.042) or Sex (*F*(1,11) = 0.660, *p* = .434, ƞ*_p_*² = 0.057), nor a significant Group by Trial interaction effect (*F*(4,44) = 1.105, *p* = .366, ƞ*_p_*² = 0.091). This suggest that all rats similarly acquired a CS-US association during the fear conditioning session, before drug infusions and avoidance training took place (Suppl. Fig. 1).

### Avoidance training

Freezing during avoidance training was scored during CS 1 and all even CS presentations (i.e., CS 2, CS 4, …, CS 30) and analyzed using a mixed ANOVA with repeated-measures factor Trial and between-subjects factors Group and Sex. The main results are reported here and the extended list of statistical results can be found in Supplementary Table 3. For an overview of the mean freezing for each avoidance training session displayed for each sex separately, see Supplementary Figure 3.

**Supplementary Figure 2.**
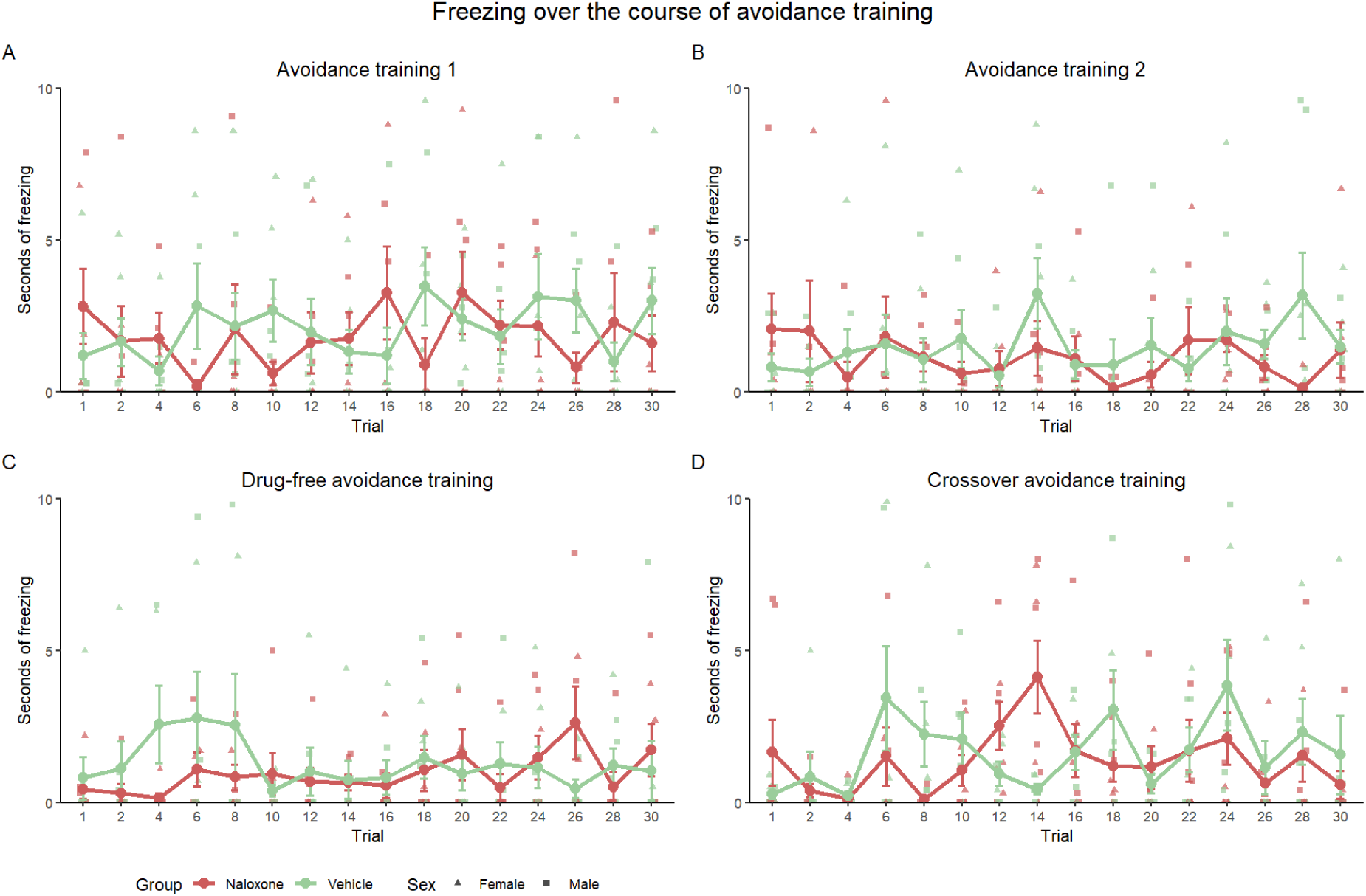
Mean ± SEM freezing for each even CS presentation (and CS 1) for rats of the naloxone (n = 7) and vehicle (n = 8 groups) for (A) avoidance training session 1, (B) avoidance training session 2, (C) the drug-free avoidance training session, and (E) the crossover avoidance training session. Triangles depict individual datapoints for female subjects and squares depict individual datapoints for male subjects.

#### Avoidance training session 1

When evaluating freezing across the first avoidance training session, we observed no significant effects of Group (*F*(1,11) = 0.205, *p* = .660, ƞ*_p_*² = 0.018), Sex (*F*(1,11) = 0.769, *p* = .399, ƞ*_p_*² = 0.065), or Trial (*F*(15,165) = 0.284, *p* = .996, ƞ*_p_*² = 0.025). Moreover, there was no significant Group by Trial interaction effect (*F*(15,165) = 1.104, *p* = .356, ƞ*_p_*² = 0.091; Suppl. Fig. 2A).

#### Avoidance training session 2

When evaluating freezing across the second avoidance training session, we observed no significant effects of Group (*F*(1,11) = 0.573, *p* = .465, ƞ*_p_*² = 0.050) or Sex (*F*(1,11) = 0.068, *p* = .800, ƞ*_p_*² = 0.006). There was a significant effect of Trial (*F*(15,165) = 1.773, *p* = **.042**, ƞ*_p_*² = 0.139), illustrating that freezing decreased across the session. Finally, there was no significant Group by Trial interaction effect (*F*(15,165) = 0.582, *p* = .886, ƞ*_p_*² = 0.050; Suppl. Fig. 2B).

#### Drug-free avoidance training session

When evaluating freezing across the drug-free avoidance training session, we observed no significant effects of Group (*F*(1,11) = 1.810, *p* = .206, ƞ*_p_*² = 0.141), Sex (*F*(1,11) = 0.019, *p* = .892, ƞ*_p_*² = 0.002) or Trial (*F*(15,165) = 1.083, *p* = .376, ƞ*_p_*² = 0.090). Finally, there was no significant Group by Trial interaction effect (*F*(15,165) = 1.439, *p* = .135, ƞ*_p_*² = 0.116; Suppl. Fig. 2C).

#### Crossover avoidance training session

When evaluating freezing across the crossover avoidance training session, we observed no significant effect of Group (*F*(1,11) = 0.230, *p* = .641, ƞ*_p_*² = 0.020), Sex (*F*(1,11) = 0.641, *p* = .440, ƞ*_p_*² = 0.055), or Trial (*F*(15,165) = 0.602, *p* = .871, ƞ*_p_*² = 0.052). Finally, there was no significant Group by Trial interaction effect (*F*(15,165) = 0.947, *p* = .514, ƞ*_p_*² = 0.079; Suppl. Fig. 2D).

**Supplementary Figure 3.**
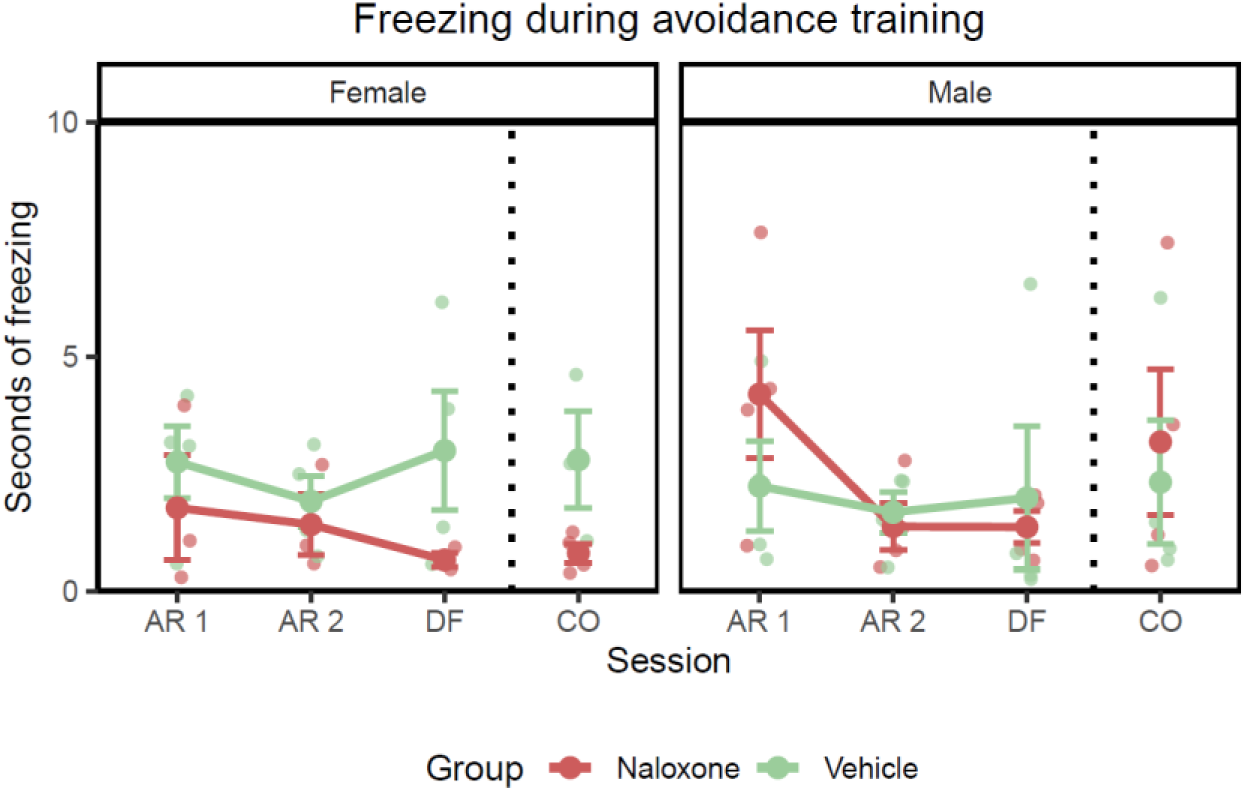
Mean ± SEM freezing for rats that received naloxone (n = 7) or vehicle (n = 8) for each avoidance training session.

### Freezing during 5-minute habituation

The main results are reported here and the extended list of statistical results can be found in Supplementary Table 4.

#### Avoidance training session 1

There was no significant effect of Minute (*F*(1.61,17.75) = 1.045, *p* = .358, ƞ*_p_*² = 0.087), nor of Group (*F*(1,11) = 0.081, *p* = .781, ƞ*_p_*² = 0.007) and Sex (*F*(1,11) = 0.045, *p* = .837, ƞ*_p_*² = 0.004). There was no significant Minute by Group interaction effect (*F*(1.61,17.75) = 0.720, *p* = .472, ƞ*_p_*² = 0.061). Interestingly, there was a significant Group by Sex interaction effect (*F*(1,11) = 9.735, *p* = **.010**, ƞ*_p_*² = 0.469), but the individual contrasts did not survive correction for multiple testing (*p*s > .193).

#### Avoidance training session 2

We did not observe significant effects of Minute (*F*(1.69,18.55) = 2.226, *p* = .142, ƞ*_p_*² = 0.168), Group (*F*(1,11) = 0.107, *p* = .848, ƞ*_p_*² = 0.010) or Sex (*F*(1,11) = 0.200, *p* = .663, ƞ*_p_*² = 0.018). There was a marginally nonsignificant Group by Minute interaction effect (*F*(1.69,18.55) = 3.520, *p* = .057, ƞ*_p_*² = 0.242), with rats of the vehicle group showing a tendency to increased freezing during the last minute of habituation (comparisons: Min 1 – Min 5: *t_Tukey_* = -3.747, *p* = **.023**; Min 2 – Min 5: *t_Tukey_* = -3.805, *p* = **.020**: Min 3 – Min 5: *t_Tukey_* = -3.646, *p* = **.030**).

#### Drug-free avoidance training session

We did not observe significant effects of Minute (*F*(1.05,11.58) = 1.201, *p* = .299, ƞ*_p_*² = 0.098), Group (*F*(1,11) = 0.454, *p* = .514, ƞ*_p_*² = 0.040), or Sex (*F*(1,11) = 1.202, *p* = .296, ƞ*_p_*² = 0.044). There was also no significant Group by Minute interaction effect (*F*(1.05,11.58) = 0.536, *p* = .488, ƞ*_p_*² = 0.046).

Crossover avoidance training session. We observed a significant effect of Minute (*F*(2.21,24.30) = 4.162, *p* = **.025**, ƞ*_p_*² = 0.274), but no significant effects of Group (*F*(1,11) = 1.585, *p* = .234, ƞ*_p_*² = 0.004) or Sex (*F*(1,11) = 0.710, *p* = .417, ƞ*_p_*² = 0.061). There was no significant interaction between Group and Minute (*F*(2.21,24.30) = 0.508, *p* = .626, ƞ*_p_*² = 0.044).

## Latency to avoid and escape the US

### Latency to avoid the US

#### Avoidance training session 1

During the first avoidance training session, we observed no significant effects of Group (*F*(1,11) = 0.118, *p* = .737, ƞ*_p_*² = 0.011), Sex (*F*(1,11) = 0.193, *p* = .669, ƞ*_p_*² = 0.017), and no significant Group by Sex interaction (*F*(1,11) = 0.023, *p* = .881, ƞ*_p_*² = 0.002), suggesting that both groups have similar latencies to avoid the US (Suppl. Fig. 4).

#### Avoidance training session 2

We observed no significant effects of Group (*F*(1,11) = 0.269, *p* = .614, ƞ*_p_*² = 0.024), Sex (*F*(1,11) = 0.241, *p* = .633, ƞ*_p_*² = 0.021), nor a significant Group by Sex interaction effect (*F*(1,11) = 1.113, *p* = .314, ƞ*_p_*² = 0.092; Suppl. Fig. 4).

#### Drug-free avoidance training session

We observed no significant effects of Group (*F*(1,11) = 0.076, *p* = .788, ƞ*_p_*² = 0.007). In line with the avoidance data, we did observe a marginally nonsignificant effect of Sex (*F*(1,11) = 4.337, *p* = .061, ƞ*_p_*² = 0.283), with females showing a tendency to avoid the US faster than males. We observed no significant Group by Sex interaction effect (*F*(1,11) = 1.346, *p* = .271, ƞ*_p_*² = 0.109; Suppl. Fig. 4).

#### Crossover avoidance training session

For the crossover avoidance training session, we observed a marginally nonsignificant main effect of Group (*F*(1,11) = 3.642, *p* = .083, ƞ*_p_*² = 0.249), as well as a marginally nonsignificant Group by Sex interaction effect (*F*(1,11) = 4.84, *p* = .05, ƞ*_p_*² = 0.306). The main effect of Sex was not significant (*F*(1,11) = 1.303, *p* = .278, ƞ*_p_*² = 0.106; Suppl. Fig. 4).

**Supplementary Figure 4.**
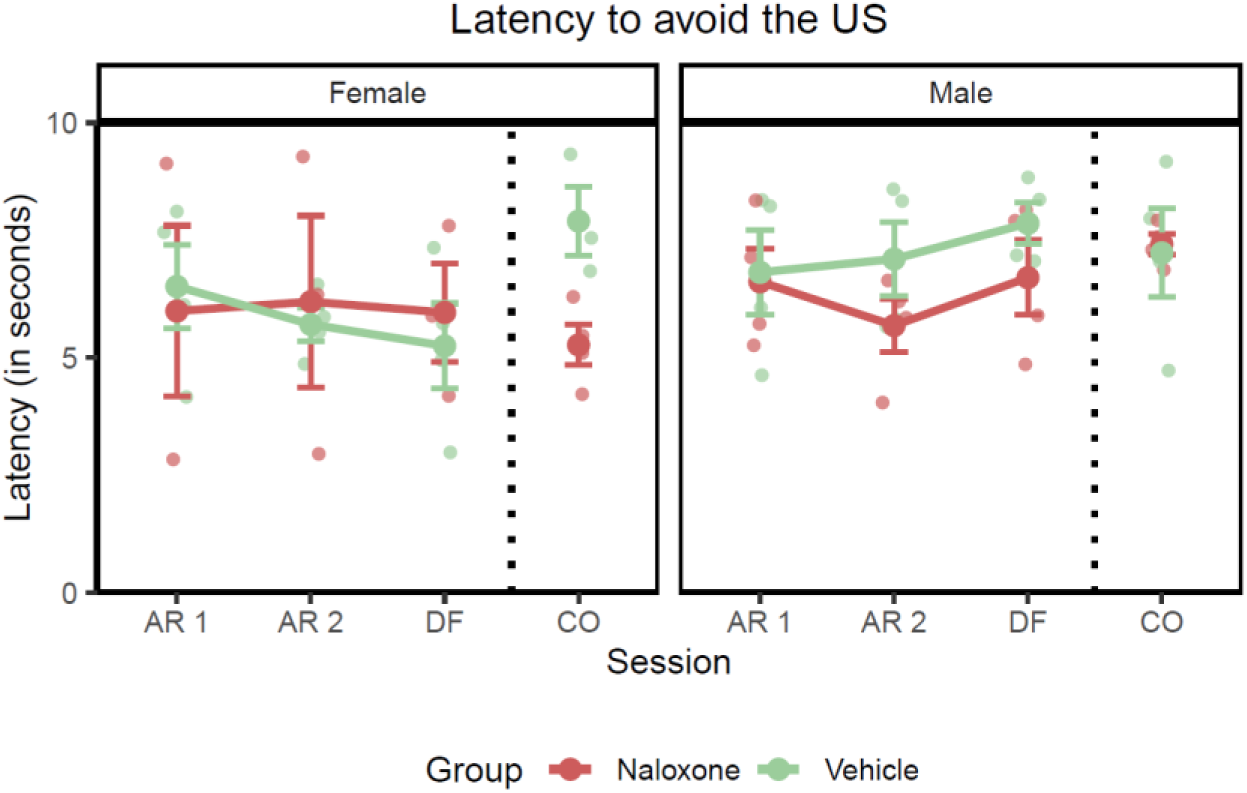
Mean ± SEM latency to avoid the US during each training session of the two-way active avoidance task for rats of the naloxone (n = 7) and vehicle (n = 8) groups. Results are plotted for males and females separately.

### Latency to escape the US

#### Avoidance training session 1

During the first avoidance training session, we observed no significant effect of Group (*F*(1,11) = 0.141, *p* = .715, ƞ*_p_*² = 0.013). The main effect of Sex was marginally nonsignificant (*F*(1,11) = 4.644, *p* = .054, ƞ*_p_*² = 0.297). There was no significant Group by Sex interaction effect (*F*(1,11) = 0.011, *p* = .919, ƞ*_p_*² < 0.001), suggesting that naloxone did not increase the latency to escape the foot shock (Suppl. Fig. 5).

#### Avoidance training session 2

During the second avoidance training session, one subject was excluded from the analyses because this subject performed 30 out of 30 avoidance responses, and thus did not receive the US during this session. We observed no significant effect of Group (*F*(1,10) = 3.527, *p* = .090, ƞ*_p_*² = 0.261), although there was a small (nonsignificant) tendency for rats that received naloxone to escape the foot shock faster than vehicle rats. The main effect of Sex was not significant (*F*(1,10) = 1.931, *p* = .195, ƞ*_p_*² = 0.162), nor was the Group by Sex interaction effect (*F*(1,10) = 0.521, *p* = .487, ƞ*_p_*² = 0.049; Suppl. Fig. 5).

#### Drug-free avoidance training session

During the drug-free avoidance session, we observed no significant effects of Group (*F*(1,11) = 0.199, *p* = .664, ƞ*_p_*² = 0.018), Sex (*F*(1,11) = 0.194, *p* = .668, ƞ*_p_*² = 0.017), nor a Group by Sex interaction effect (*F*(1,11) = 0.245, *p* = .631, ƞ*_p_*² = 0.022; Suppl. Fig. 5).

#### Crossover avoidance training session

For the crossover avoidance training session, we observed no significant effects of Group (*F*(1,11) = 2.576, *p* = .137, ƞ*_p_*² = 0.190) and Sex (*F*(1,11) = 2.009, *p* = .184, ƞ*_p_*² = 0.154). The Group by Sex interaction effect was marginally nonsignificant (*F*(1,11) = 3.726, *p* = .080, ƞ*_p_*² = 0.253; Suppl. Fig. 5).

**Supplementary Figure 5.**
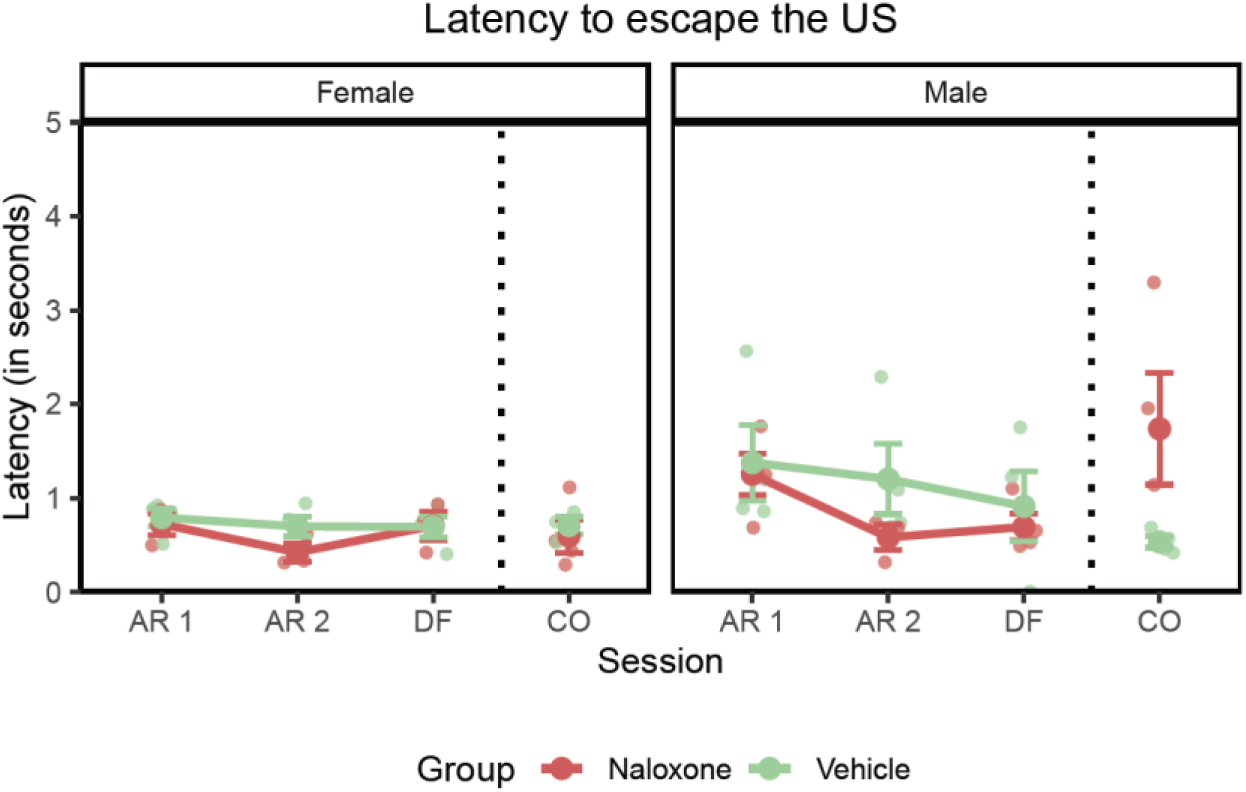
Mean ± SEM latency to escape the US during each training session of the two-way active avoidance task for rats of the naloxone (n = 7) and vehicle (n = 8) groups. Results are plotted for males and females separately.

## Number of shuttles during avoidance training

### Number of shuttles during the 5-minute habituation

#### Avoidance training session 1

We observed no significant effects of Group (*F*(1,11) = 0.913, *p* = .360, ƞ*_p_*² = 0.077), Sex (*F*(1,11) = 0.392, *p* = .544, ƞ*_p_*² = 0.034) nor a significant Group by Sex interaction effect (*F*(1,11) = 3.286, *p* = .097, ƞ*_p_*² = 0.230).

#### Avoidance training session 2

We observed no significant effects of Group (*F*(1,11) = 0.035, *p* = .856, ƞ*_p_*² = 0.003), Sex (*F*(1,11) = 0.553, *p* = .473, ƞ*_p_*² = 0.167), nor a significant Group by Sex interaction (*F*(1,11) = 2.210, *p* = .165, ƞ*_p_*² = 0.167).

#### Drug-free avoidance training session

We observed no significant effects of Group (*F*(1,11) = 0.057, *p* = .816, ƞ*_p_*² = 0.005), Sex (*F*(1,11) = 0.228, *p* = .642, ƞ*_p_*² = 0.115), nor a significant Group by Sex interaction (*F*(1,11) = 1.426, *p* = .258, ƞ*_p_*² = 0.115).

#### Crossover avoidance training session

We observed no significant effects of Group (*F*(1,11) = 0.261, *p* = .620, ƞ*_p_*² = 0.023), Sex (*F*(1,11) = 0.083, *p* = .778, ƞ*_p_*² = 0.008), nor a significant Group by Sex interaction (*F*(1,11) = 1.836, *p* = .203, ƞ*_p_*² = 0.143).

### Number of shuttles during ITIs

#### Avoidance training session 1

We observed no significant effects of Group (*F*(1,11) = 2.875, *p* = .118, ƞ*_p_*² = 0.207) and Sex (*F*(1,11) = 0.939, *p* = .353, ƞ*_p_*² = 0.079). Interestingly, there was a significant Group by Sex interaction effect (*F*(1,11) = 6.972, *p* = **.023**, ƞ*_p_*² = 0.388), but the individual contrasts did not survive correction for multiple testing.

#### Avoidance training session 2

We observed no significant effects of Group (*F*(1,11) = 1.682, *p* = .221, ƞ*_p_*² = 0.122), Sex (*F*(1,11) = 1.868, *p* = .199, ƞ*_p_*² = 0.145), nor a significant Group by Sex interaction effect (*F*(1,11) = 3.188, *p* = .102, ƞ*_p_*² = 0.225).

#### Drug-free avoidance training session

We observed no significant effects of Group (F(1,11) = 0.031, *p* = .864, ƞ*_p_*² = 0.003), Sex (*F*(1,11) = 1.350, *p* = .270, ƞ*_p_*² = 0.109), nor a significant Group by Sex interaction effect (*F*(1,11) = 0.995, *p* = .340, ƞ*_p_*² = 0.083).

#### Crossover avoidance training session

We observed a marginally nonsignificant effect of Group (*F*(1,11) = 3.289, *p* = .097, ƞ*_p_*² = 0.230), with a tendency for naloxone to increase shuttling. The main effect of Sex was also marginally nonsignificant (*F*(1,11) = 4.247, *p* = .0.064, ƞ*_p_*² = 0.279), with a tendency for females to shuttle more than males. The Group by Sex interaction was not significant (*F*(1,11) = 0.322, *p* = .582, ƞ*_p_*² = 0.028).

## Number of escape failures

We exploratively examined the number of escape failures to assess whether naloxone increases the number of escape failures. We observed no escape failures during avoidance training session 1 and avoidance training session 2. One rat showed 3 escape failures during the drug-free test and 2 escape failures during the crossover test (when it received vehicle), suggesting that naloxone did not increase the number of escape failures.

## Number of avoidance responses in blocks, per sex

**Supplementary Figure 6.**
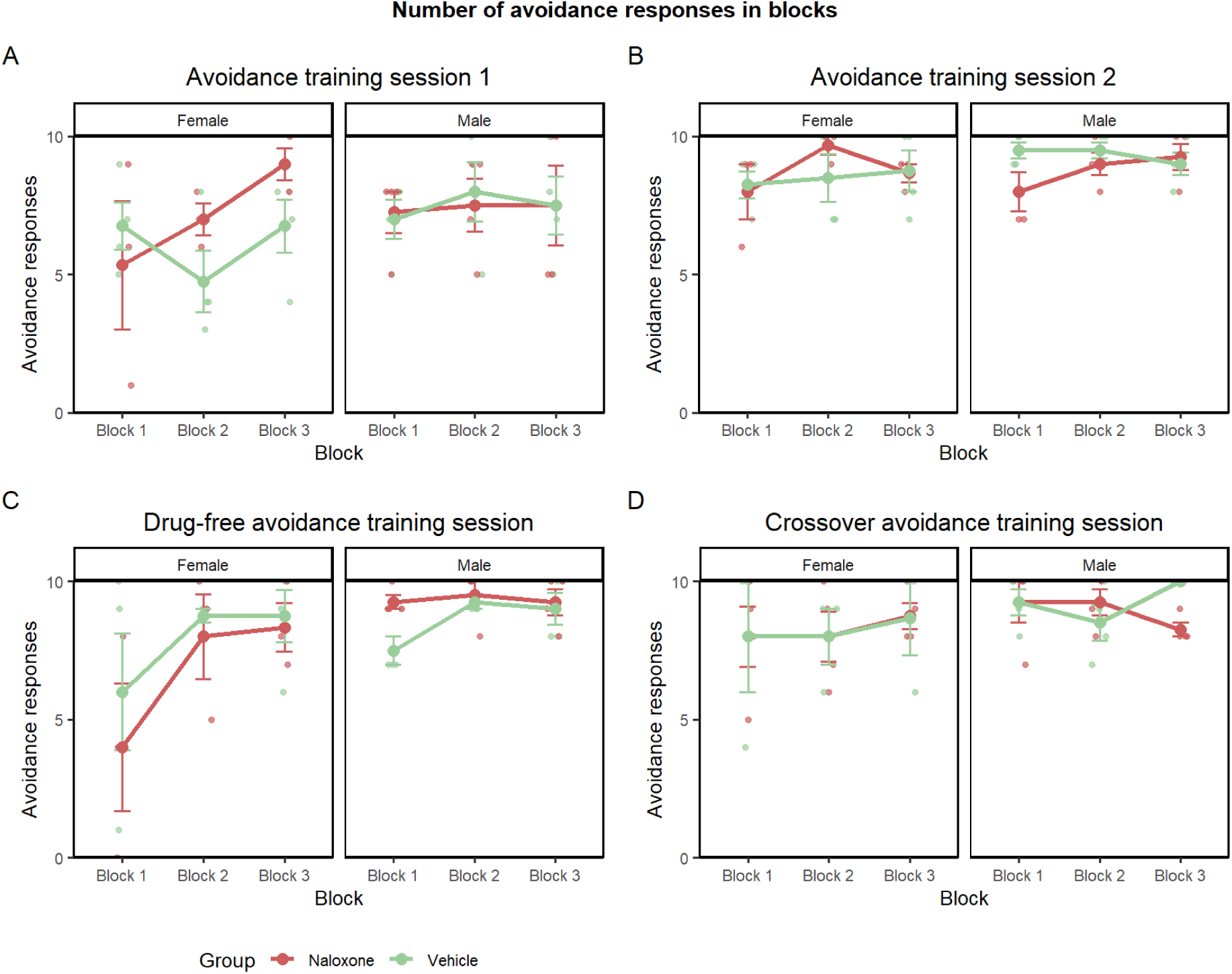
Mean (± SEM) number of avoidance responses in three blocks of 10 trials each for the naloxone (n = 7) and vehicle (n = 8) groups, displayed separately for males and females, for (A) avoidance training session 1, (B) avoidance training session 2, (C) the drug-free avoidance training session, and (D) the crossover avoidance training session.

## Additional analyses: post-shock freezing and grooming

### Post-shock freezing after non-avoided shocks during the first block of avoidance training 1

We conducted a non-parametric test because the Shapiro-Wilk test indicated a deviation from normality (p = .007). A one-sided Mann-Whitney test found no significant difference between the naloxone and vehicle groups (U = 23.00, p = .418, effect size (biserial rank correlation) = -0.095).

**Supplementary Figure 7.**
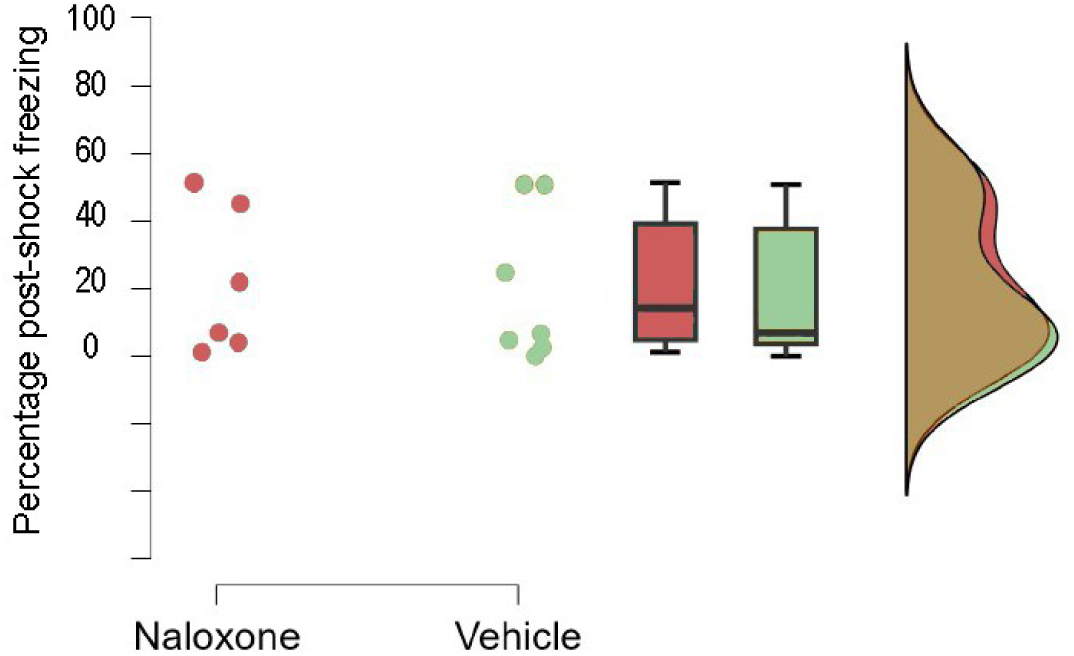
Raincloud plot showing the naloxone (n = 6) and vehicle (n = 7) groups. Individual observations are shown as dots, boxplots indicate the median and interquartile range, and additional plots represent data density.

### Post-shock grooming after non-avoided shocks during the first block of avoidance training 1

We observed no grooming in any of the animals during this period.

### Grooming during the 5-min acclimation period of avoidance training 1

A one-sided t-test found no significant difference between the naloxone and vehicle groups (t = 1.164, p = .865, Cohen’s d = 0.647).

**Supplementary Figure 8.**
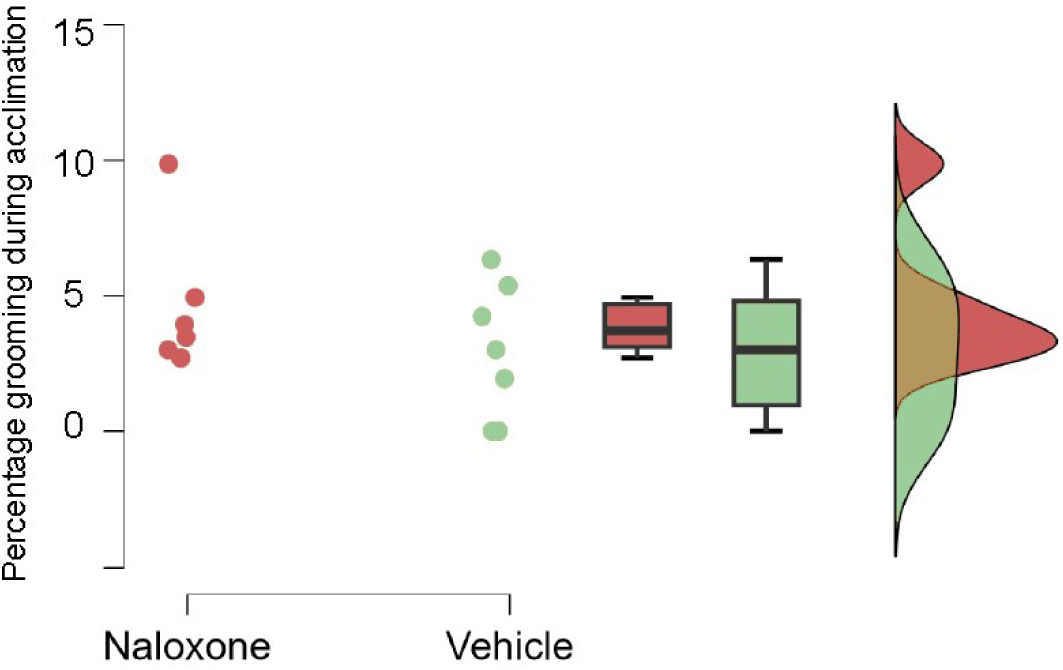
Raincloud plot showing the naloxone (n = 6) and vehicle (n = 7) groups. Individual observations are shown as dots, boxplots indicate the median and interquartile range, and additional plots represent data density.

## Supplementary Tables

**Supplementary Table 1.**
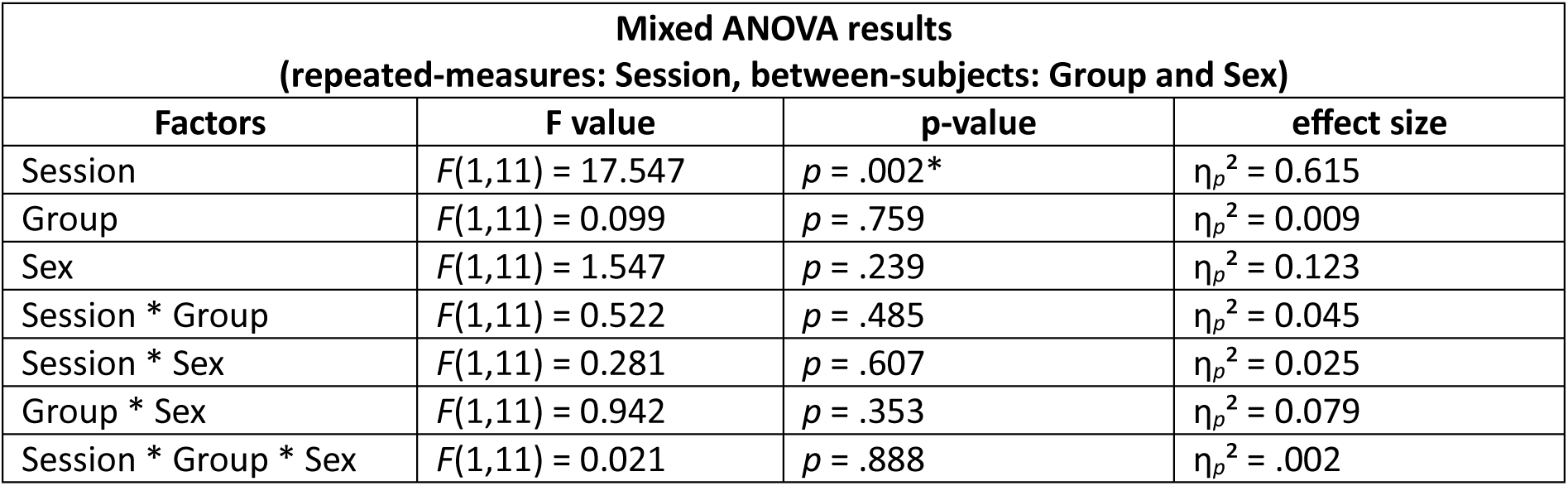
Overview of the statistical results of a mixed ANOVA with repeated-measures factor Session (2 levels: Session 1 and Session 2) and between-subjects factors Group (naloxone and vehicle) and Sex (male and female). Significant effects are indicated by * (*p* < .05).

| Mixed ANOVA results<br>(repeated-measures: Session, between-subjects: Group and Sex) |  |  |  |
| --- | --- | --- | --- |
| Factors | F value | p-value | effect size |
| Session | $F(1,11) = 17.547$ | $p = .002^*$ | $\eta_p^2 = 0.615$ |
| Group | $F(1,11) = 0.099$ | $p = .759$ | $\eta_p^2 = 0.009$ |
| Sex | $F(1,11) = 1.547$ | $p = .239$ | $\eta_p^2 = 0.123$ |
| Session * Group | $F(1,11) = 0.522$ | $p = .485$ | $\eta_p^2 = 0.045$ |
| Session * Sex | $F(1,11) = 0.281$ | $p = .607$ | $\eta_p^2 = 0.025$ |
| Group * Sex | $F(1,11) = 0.942$ | $p = .353$ | $\eta_p^2 = 0.079$ |
| Session * Group * Sex | $F(1,11) = 0.021$ | $p = .888$ | $\eta_p^2 = .002$ |

**Supplementary Table 2.** Overview of the statistical results of a mixed ANOVA with repeated-measures factor Block (3 levels: Block 1: first 10 trials, Block 2: middle 10 trials, Block 3: final 10 trials) and between-subjects factors Group (naloxone and vehicle) and Sex (male and female) for each avoidance training session. Significant effects are indicated by * (*p* < .05).

| **Mixed ANOVA results**<br>**(repeated-measures: Block, between-subjects: Group and Sex)** |  |  |  |
| --- | --- | --- | --- |
| **Avoidance training session 1** |  |  |  |
| **Factors** | **F value** | **p-value** | **effect size** |
| Block | $F(1.34, 14.68) = 2.084$ | $p = .168$ | $\eta_p^2 = 0.159$ |
| Group | $F(1, 11) = 0.290$ | $p = .601$ | $\eta_p^2 = 0.026$ |
| Sex | $F(1, 11) = 0.964$ | $p = .347$ | $\eta_p^2 = 0.081$ |
| Block * Group | $F(1.34, 14.68) = 1.306$ | $p = .286$ | $\eta_p^2 = 0.106$ |
| Block * Sex | $F(1.34, 14.68) = 1.999$ | $p = .178$ | $\eta_p^2 = 0.154$ |
| Group * Sex | $F(1, 11) = 0.401$ | $p = .539$ | $\eta_p^2 = 0.035$ |
| Block * Group * Sex | $F(1.34, 14.68) = 2.244$ | $p = .152$ | $\eta_p^2 = 0.169$ |
| **Avoidance training session 2** |  |  |  |
| Block | $F(2, 22) = 1.731$ | $p = .200$ | $\eta_p^2 = 0.136$ |
| Group | $F(1, 11) = 0.192$ | $p = .670$ | $\eta_p^2 = 0.017$ |
| Sex | $F(1, 11) = 1.335$ | $p = .272$ | $\eta_p^2 = 0.108$ |
| Block * Group | $F(2, 22) = 1.282$ | $p = .297$ | $\eta_p^2 = 0.104$ |
| Block * Sex | $F(2, 22) = 0.166$ | $p = .848$ | $\eta_p^2 = 0.015$ |
| Group * Sex | $F(1, 11) = 1.526$ | $p = .243$ | $\eta_p^2 = 0.122$ |
| Block * Group * Sex | $F(2, 22) = 0.877$ | $p = .430$ | $\eta_p^2 = 0.074$ |
| **Drug-free avoidance training session** |  |  |  |
| Block | $F(1.29, 14.17) = 5.246$ | $p = .031^*$ | $\eta_p^2 = 0.323$ |
| Group | $F(1, 11) = 0.085$ | $p = .776$ | $\eta_p^2 = 0.008$ |
| Sex | $F(1, 11) = 9.944$ | $p = .009^*$ | $\eta_p^2 = 0.475$ |
| Block * Group | $F(1.29, 14.17) = 0.006$ | $p = .969$ | $\eta_p^2 < 0.001$ |
| Block * Sex | $F(1.29, 14.17) = 1.900$ | $p = .190$ | $\eta_p^2 = 0.147$ |
| Group * Sex | $F(1, 11) = 2.967$ | $p = .113$ | $\eta_p^2 = 0.212$ |
| Block * Group * Sex | $F(1.29, 14.17) = 0.600$ | $p = .492$ | $\eta_p^2 = 0.052$ |
| **Crossover avoidance training session** |  |  |  |
| Block | $F(2, 22) = 0.897$ | $p = .422$ | $\eta_p^2 = 0.075$ |
| Group | $F(1, 11) = 0.044$ | $p = .837$ | $\eta_p^2 = 0.004$ |
| Sex | $F(1, 11) = 1.360$ | $p = .268$ | $\eta_p^2 = 0.110$ |
| Block * Group | $F(2, 22) = 1.472$ | $p = .251$ | $\eta_p^2 = 0.118$ |
| Block * Sex | $F(2, 22) = 0.670$ | $p = .522$ | $\eta_p^2 = 0.057$ |
| Group * Sex | $F(1, 11) = 0.062$ | $p = .808$ | $\eta_p^2 = 0.006$ |
| Block * Group * Sex | $F(2, 22) = 1.699$ | $p = .206$ | $\eta_p^2 = 0.134$ |

**Supplementary Table 3.** Statistical results of the ANOVAs conducted for freezing during all trials of the Pavlovian fear conditioning session and freezing during the first CS and all even CS presentations of each avoidance training session. Significant effects are indicated with * (*p* < .05).

| **Mixed ANOVA on freezing**<br>**(repeated-measures: Trial, between-subjects: Group and Sex)** |  |  |  |
| --- | --- | --- | --- |
| **Pavlovian fear conditioning session** |  |  |  |
| **Factors** | **F value** | **p-value** | **effect size** |
| Trial | $F(4,44) = 10.034$ | $p < .001^*$ | $\eta_p^2 = 0.477$ |
| Group | $F(1,11) = 0.478$ | $p = .504$ | $\eta_p^2 = 0.042$ |
| Sex | $F(1,11) = 0.660$ | $p = .434$ | $\eta_p^2 = 0.057$ |
| Trial * Group | $F(4,44) = 1.105$ | $p = .366$ | $\eta_p^2 = 0.091$ |
| Trial * Sex | $F(4,44) = 1.168$ | $p = .338$ | $\eta_p^2 = 0.096$ |
| Group * Sex | $F(1,11) = 0.001$ | $p = .972$ | $\eta_p^2 < .001$ |
| Trial * Group * Sex | $F(4,44) = 0.136$ | $p = .968$ | $\eta_p^2 = 0.012$ |
| **Avoidance training session 1** |  |  |  |
| Trial | $F(15,165) = 0.284$ | $p = .996$ | $\eta_p^2 = 0.025$ |
| Group | $F(1,11) = 0.205$ | $p = .660$ | $\eta_p^2 = 0.018$ |
| Sex | $F(1,11) = 0.769$ | $p = .399$ | $\eta_p^2 = 0.065$ |
| Trial * Group | $F(15,165) = 1.104$ | $p = .356$ | $\eta_p^2 = 0.091$ |
| Trial * Sex | $F(15,165) = 0.735$ | $p = .747$ | $\eta_p^2 = 0.063$ |
| Group * Sex | $F(1,11) = 1.831$ | $p = .203$ | $\eta_p^2 = 0.143$ |
| Trial * Group * Sex | $F(15,165) = 0.942$ | $p = .519$ | $\eta_p^2 = 0.079$ |
| **Avoidance training session 2** |  |  |  |
| Trial | $F(15,165) = 1.773$ | $p = .042^*$ | $\eta_p^2 = 0.139$ |
| Group | $F(1,11) = 0.573$ | $p = .465$ | $\eta_p^2 = 0.050$ |
| Sex | $F(1,11) = 0.068$ | $p = .800$ | $\eta_p^2 = 0.006$ |
| Trial * Group | $F(15,165) = 0.582$ | $p = .886$ | $\eta_p^2 = 0.050$ |
| Trial * Sex | $F(15,165) = 1.276$ | $p = .223$ | $\eta_p^2 = 0.104$ |
| Group * Sex | $F(1,11) = 0.033$ | $p = .860$ | $\eta_p^2 = 0.003$ |
| Trial * Group * Sex | $F(15,165) = 0.546$ | $p = .911$ | $\eta_p^2 = 0.047$ |
| **Drug-free avoidance training session** |  |  |  |
| Trial | $F(15,165) = 1.083$ | $p = .376$ | $\eta_p^2 = 0.090$ |
| Group | $F(1,11) = 1.810$ | $p = .206$ | $\eta_p^2 = 0.141$ |
| Sex | $F(1,11) = 0.019$ | $p = .892$ | $\eta_p^2 = 0.002$ |
| Trial * Group | $F(15,165) = 1.439$ | $p = .135$ | $\eta_p^2 = 0.116$ |
| Trial * Sex | $F(15,165) = 1.467$ | $p = .123$ | $\eta_p^2 = 0.118$ |
| Group * Sex | $F(1,11) = 0.609$ | $p = .451$ | $\eta_p^2 = 0.052$ |
| Trial * Group * Sex | $F(15,165) = 1.385$ | $p = .160$ | $\eta_p^2 = 0.112$ |
| **Crossover avoidance training session** |  |  |  |
| Trial | $F(15,165) = 0.602$ | $p = .871$ | $\eta_p^2 = 0.052$ |
| Group | $F(1,11) = 0.230$ | $p = .641$ | $\eta_p^2 = 0.020$ |
| Sex | $F(1,11) = 0.641$ | $p = .440$ | $\eta_p^2 = 0.055$ |
| Trial * Group | $F(15,165) = 0.947$ | $p = .514$ | $\eta_p^2 = 0.079$ |
| Trial * Sex | $F(15,165) = 1.099$ | $p = .361$ | $\eta_p^2 = 0.091$ |
| Group * Sex | $F(1,11) = 1.452$ | $p = .253$ | $\eta_p^2 = 0.117$ |
| Trial * Group * Sex | $F(15,165) = 0.764$ | $p = .716$ | $\eta_p^2 = 0.065$ |

**Supplementary Table 4.** Freezing during the first five minutes of habituation of each avoidance training session, scored per minute and analyzed using a mixed ANOVA with repeated-measures factor Minute and between-subjects factors Group and Sex. Significant effects are indicated by * (*p* < .05).

| **Mixed ANOVA on freezing**<br>**(repeated-measures: Minute, between-subjects: Group and Sex)** |  |  |  |
| --- | --- | --- | --- |
| **Habituation of avoidance training session 1** |  |  |  |
| **Factors** | **F value** | **p-value** | **effect size** |
| Minute | $F(1.61,17.75) = 1.045$ | $p = .358$ | $\eta_p^2 = 0.087$ |
| Group | $F(1,11) = 0.081$ | $p = .781$ | $\eta_p^2 = 0.007$ |
| Sex | $F(1,11) = 0.045$ | $p = .837$ | $\eta_p^2 = 0.004$ |
| Minute * Group | $F(1.61,17.75) = 0.720$ | $p = .472$ | $\eta_p^2 = 0.061$ |
| Minute * Sex | $F(1.61,17.75) = 0.866$ | $p = .416$ | $\eta_p^2 = 0.073$ |
| Group * Sex | $F(1,11) = 9.735$ | $p = .010^*$ | $\eta_p^2 = 0.469$ |
| Minute * Group * Sex | $F(1.61,17.75) = 0.711$ | $p = .476$ | $\eta_p^2 = 0.061$ |
| **Habituation avoidance training session 2** |  |  |  |
| Minute | $F(1.69,18.55) = 2.226$ | $p = .142$ | $\eta_p^2 = 0.168$ |
| Group | $F(1,11) = 0.107$ | $p = .750$ | $\eta_p^2 = 0.010$ |
| Sex | $F(1,11) = 0.200$ | $p = .663$ | $\eta_p^2 = 0.018$ |
| Minute * Group | $F(1.69,18.55) = 3.520$ | $p = .057$ | $\eta_p^2 = 0.242$ |
| Minute * Sex | $F(1.69,18.55) = 0.973$ | $p = .383$ | $\eta_p^2 = 0.081$ |
| Group * Sex | $F(1,11) = 0.038$ | $p = .848$ | $\eta_p^2 = 0.003$ |
| Minute * Group * Sex | $F(1.69,18.55) = 0.590$ | $p = .537$ | $\eta_p^2 = 0.051$ |
| **Habituation drug-free avoidance training session** |  |  |  |
| Minute | $F(1.05,11.58) = 1.201$ | $p = .299$ | $\eta_p^2 = 0.098$ |
| Group | $F(1,11) = 0.454$ | $p = .514$ | $\eta_p^2 = 0.040$ |
| Sex | $F(1,11) = 1.202$ | $p = .296$ | $\eta_p^2 = 0.099$ |
| Minute * Group | $F(1.05,11.58) = 0.536$ | $p = .488$ | $\eta_p^2 = 0.046$ |
| Minute * Sex | $F(1.05,11.58) = 1.030$ | $p = .336$ | $\eta_p^2 = 0.086$ |
| Group * Sex | $F(1,11) = 0.509$ | $p = .491$ | $\eta_p^2 = 0.044$ |
| Minute * Group * Sex | $F(1.05,11.58) = 0.565$ | $p = .476$ | $\eta_p^2 = 0.049$ |
| **Habituation crossover avoidance training session** |  |  |  |
| Minute | $F(2.21,24.30) = 4.162$ | $p = .025^*$ | $\eta_p^2 = 0.274$ |
| Group | $F(1,11) = 1.585$ | $p = .234$ | $\eta_p^2 = 0.126$ |
| Sex | $F(1,11) = 0.710$ | $p = .417$ | $\eta_p^2 = 0.061$ |
| Minute * Group | $F(2.21,24.30) = 0.508$ | $p = .626$ | $\eta_p^2 = 0.044$ |
| Minute * Sex | $F(2.21,24.30) = 0.415$ | $p = .684$ | $\eta_p^2 = 0.036$ |
| Group * Sex | $F(1,11) = 0.043$ | $p = .840$ | $\eta_p^2 = 0.004$ |
| Minute * Group * Sex | $F(2.21,24.30) = 1.527$ | $p = .237$ | $\eta_p^2 = 0.122$ |

